# DNA co-methylation has a stable structure and is related to specific aspects of genome regulation

**DOI:** 10.1101/2022.03.16.484648

**Authors:** Sarah H Watkins, Matthew Suderman, Gibran Hemani, Kimberly Burrows, Deborah A. Lawlor, Jane West, Kathryn Willan, Nicholas J Timpson, Josine L Min, Tom R Gaunt

## Abstract

DNA methylation (DNAm) is influenced by genetic and environmental factors, and can be used to understand interindividual variability in genomic regulation. Co-methylation between DNAm sites is a known phenomenon, but the architecture of relationships between the approximately 450,000 (450k) sites commonly measured in epidemiological studies has not been described. We investigate whether interindividual co-methylation structure amongst the 450k sites changes with age, whether it differs between UK-born White (n=849, 910, 921 and 424) and Pakistani ancestry (n=439) individuals, and how it relates to genome regulation.

We find stability between birth and adolescence, across cohorts, and between two ethnic groups. Highly correlated DNAm sites in close proximity are heritable, but these relationships are weakly influenced by nearby genetic variants, and are enriched for transcription factor (TF) binding sites related to regulation of short RNAs transcribed by RNA polymerase III. Highly correlated sites that are distant, or on different chromosomes (in *trans*), are driven by common and unique environmental factors, with methylation at these sites less likely to be driven by genotype. *Trans* co-methylated DNAm sites are enriched for multiple TF binding sites and for inter-chromosomal chromatin contact sites, suggesting DNA co-methylation of distant sites may relate to long-range cooperative TF interactions.

We conclude that DNA co-methylation has a stable structure from birth to adolescence, and between UK-born White and Pakistani individuals. This stable structure might have implications for future design and interpretation of epigenetic studies. We hypothesise that co-methylation may have roles in genome regulation in humans, including 3D chromatin architecture.

## Introduction

DNA methylation (DNAm) is an epigenetic modification to DNA that can be influenced by both environmental and genetic factors (1–3). It has roles in a variety of genomic functions, including a complex and context-dependent alteration of gene expression (4–7); both altering and being altered by transcription factor binding (8–13); and interaction with chromatin states (14–17). Standardized DNAm assays have enabled epigenome wide association studies (EWAS), where each assayed DNAm site is tested independently for association with a variable of interest. These studies have identified DNAm sites associated with many exposures and diseases (18–23). Networks of co-methylated DNAm sites have also been described - these utilise the relationships between DNAm sites to infer regulatory pathways by which DNAm might relate to health or disease (24–27). In contrast to genetic architecture, our knowledge of relationships between these commonly assayed DNAm sites at the population level is still limited, constraining our understanding of the principles of epigenetic epidemiology underlying co-methylation between and across populations, and whether co-methylation has a role in genome regulation. In addition to this need for greater biological understanding, the co-methylation structure is an important consideration for the statistical design of EWAS and differentially methylated region (DMR) studies; no equivalent of genetic linkage disequilibrium (LD) matrices yet exists for imputation in DNAm analyses. A deeper understanding of DNAm correlation structure, and its stability within and between individuals at the population level, might help identify whether this structure can be used for tagging, pruning and imputation in EWAS in a similar way to how LD enables these approaches to be used in genome-wide association studies (GWAS).

Both whole genome bisulfite sequencing (WGBS) and array-based studies highlighted that DNA methylation forms local correlation structures, with DNAm sites within 1-2kb often having correlated methylation states (28–34); WGBS data shows that immediately adjacent sites almost always have the same methylation state (28). Correlations between close DNAm sites (referred to here as *cis* correlations) may be driven by genetic variants, as regions of highly correlating DNAm sites have been associated with nearby SNPs (30, 33); although other studies have shown that correlations between DNAm sites can be driven by environmental exposures (35). Co-methylation structure does not mirror the large blocks that linkage disequilibrium (LD) forms (30), and it appears to be consistent across ethnic groups with different genetic architecture (32); this is because co-methylation depends on DNA methyltransferases and demethylases, whereas LD is determined by demographic history, recombination and mutation rate. There have been conflicting reports of whether DNAm correlation structure is related to genomic annotations such as CpG islands (30, 34), but recent work has demonstrated that DNAm sites within 2Mb at which methylation level co-varies due to the same causal genetic variants can be used to predict contact sites for chromatin loops (36).

Co-methylation between distant DNAm sites on the same chromosome, and sites on different chromosomes (referred to here as *trans* correlations), are less well described. Correlation of a limited number of *trans*-chromosomal DNAm sites with highly variable DNAm levels showed that DNAm sites related to *HOX* genes were highly correlated, suggesting *trans*-chromosomal correlations might be related to biological pathways (35). Another recent paper using mouse tissue has shown that DNAm sites around inter- and intra-chromosomal chromatin contact points have correlated methylation states, with those within the same topologically associating domains and with the same chromatin states having more correlated methylation states (16). This suggests correlations between DNAm sites are likely to be relevant to genome regulation; however this has yet to be shown in human studies for inter-chromosomal contacts.

DNAm is a dynamic epigenetic mark, so it is unsurprising that measurements at individual sites are not always stable. Sites influenced by genotype tend to be more stable than those that are not; twin models have found that the heritability of DNAm is on average 19% (1, 37, 38), with a subsequent study showing the reliability of each of DNAm probe measurement is associated with the heritability of the probe (39). DNAm changes with age (40–42) and in response to environmental exposures such as smoking (18, 43), but environmental and genetic constraints have been shown to contribute to the stability of DNAm over the lifecourse (37). As yet there is no indication of how stable relationships between DNAm sites might be; a lack of stability might indicate co-methylation has changing functions or changing environmental influences, and would limit the utility of correlated methylation states in inferring genome regulation over different datasets. It is therefore important to assess correlation structure over time in the same individuals, and across datasets (to ensure replication). An important consideration when investigating stability is utilising datasets that include diverse social groups; DNA methylation is associated with adversity (44, 45), racial discrimination (46, 47), social inequalities (19, 48), and environmental exposures such as air pollution (49, 50). Consequently, co-methylation structure is more likely to reflect stable biological processes if observations persist across diverse social groups (51).

In this paper, we outline DNA co-methylation structure in blood across DNAm sites featuring on the widely used Illumina 450k Beadchip (450k array) which mainly covers promoters, TSS, and coding transcripts across 1.5% of the methylome (52, 53). We use data from two large UK birth cohorts - the Accessible Resource for Integrated Epigenomic studies (ARIES), is a sub study of the Avon Longitudinal Study of Parents and Children (ALSPAC), which recruited pregnant women in the South West of England with predicted delivery dates between April 1991 and December 1992; and Born in Bradford (BiB), a birth cohort recruited in a city in the North of England that recruited pregnant women between with predicted delivery dates between March 2007 and November 2010. ARIES has longitudinal DNAm data from 849, 910, and 921 White British participants, at birth (cord blood), 7 and 15-17 years, respectively; BiB recruited in a city with high levels of deprivation, where 55% of the obstetric population are South Asian (mostly Pakistani). The BiB subcohort with DNAm data is approximately evenly split between two UK-born ethnic groups, with 424 white British and 439 Pakistani participants, with DNAm measured at birth (in cord blood). In our analyses we split BiB by ethnic group (where membership of ethnic groups was obtained through maternal self-report, and we conceptualise ethnicity as a social construct that is associated with differing social and environmental exposures, and therefore potentially differing effects on the methylome). We assess the stability of the co-methylation structure between DNAm sites over time, across ARIES and BiB, and between two ethnic groups born in the same geographic area. We detail genetic and environmental influences on this correlation structure, and provide the first comprehensive analysis of strong *cis* and *trans* co-methylation between DNAm sites across the 450k array at the population level, outlining their distinct potential roles in genome regulation. We also provide a resource which can be used by the scientific community [DOI 10.5523/bris.31uze72mt042g2ticr0w6z6v8y].

## Results

### Co-methylation structure and stability, across the 450k array Overall co-methylation structure

To assess co-methylation structure, we adjusted DNAm data for age, sex, cell counts and batch effects and created a correlation matrix (see Methods, Figure 11) between all remaining possible pairs of sites on the 450k array in ARIES (n sites=394,842), and all sites on the EPIC array that also feature on the 450k array in BiB (n sites=369,796). To assess features of correlations of different strengths, co-methylated pairs were aggregated into bands of correlation, ranging from −1 to 1, in increments of 0.1 (see Methods for details). For all 5 datasets, the distributions of all possible pairwise correlations are positively skewed, and 83-87% of correlations are between −0.2 and 0.2 (Figure 1; see supplementary figure 1 for the plot split by *cis* and *trans* correlations). To investigate whether physical proximity has an influence on co-methylation, we defined *cis* as within 1Mb and *trans* as over 1Mb. On average, just 12.6% of *cis* and 14.5% of *trans* pairs had a correlation above 0.2 or below −0.2. Physical proximity between DNAm sites influences the likelihood of them having highly correlated methylation states (Table 1) - for the strongest positive correlations (R=0.9-1) we see a greater proportion of *cis* than *trans* correlations (p=<2.2e-16 in all datasets using a chi squared test). We also see a trend of stronger correlations in BiB than in ARIES - for the strongest positive correlations (R=0.9-1) we see a greater proportion of *trans* correlations in the BiB cohort than in ARIES (p=<2.2e-16 using a chi squared test at birth between ARIES and the BiB white British individuals), and for strong positive correlations (R>0.5) we see a greater proportion of both *cis* and *trans* correlations in BiB than in ARIES (p=<2.2e-16 using a chi squared test at birth between ARIES and the BiB white British individuals). However this may be due to a thresholding effect; see section “Stability of correlations across time, datasets, and ethnic groups”. As strongly co-methylated pairs (R>0.5 and R<-0.5) reflect less than 15% of all correlations, they can be viewed more clearly in Table 1 than Figure 1.

**Figure 1:**
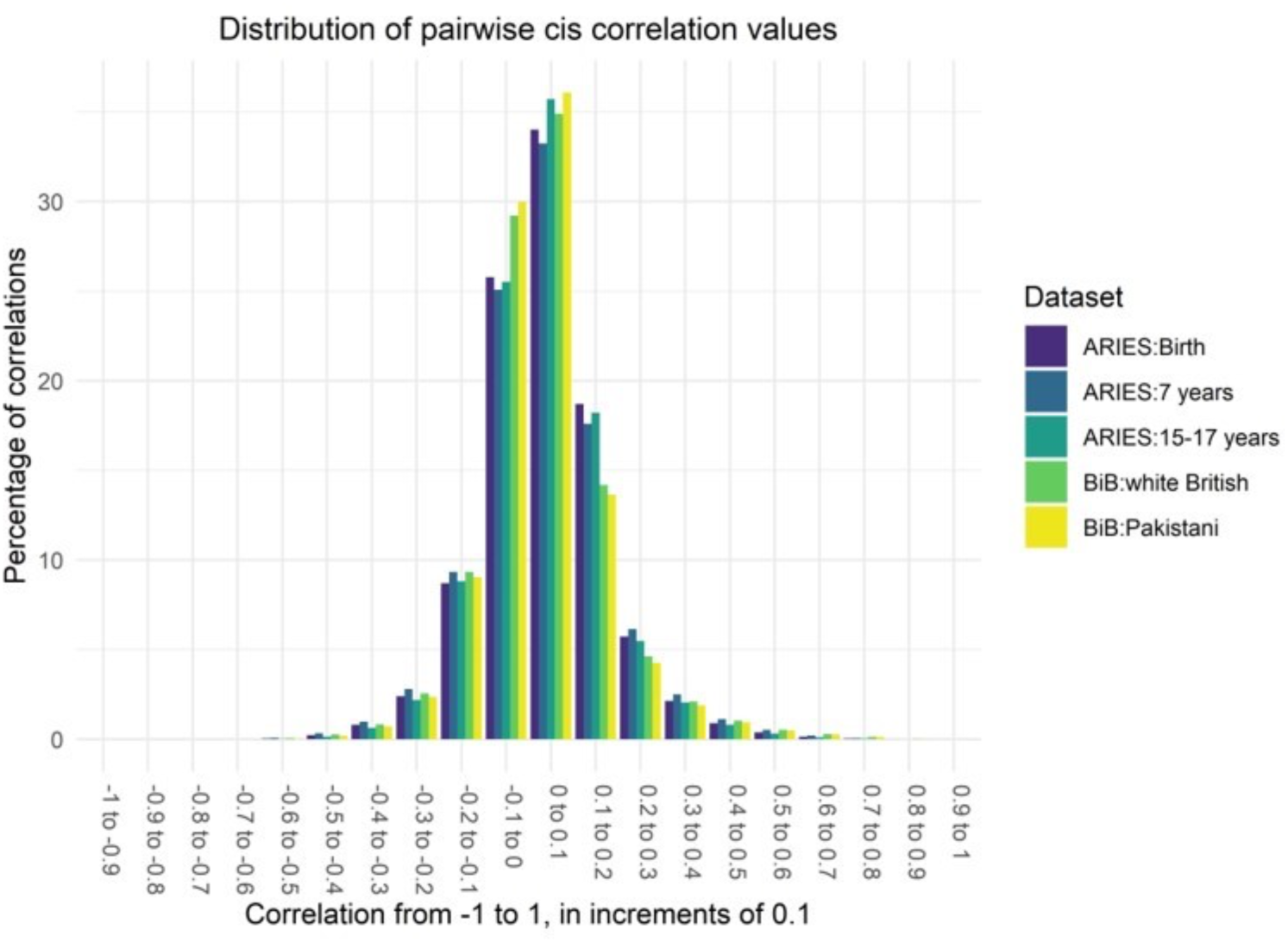
Bar plot showing the distribution of biweight mid-correlation values between all filtered DNAm sites on the 450k array (ARIES, n sites=394,842) and all filtered DNAm sites on the EPIC array that also feature on the 450k array (BiB, n sites=369,796)

**Table 1:** Table of the numbers (percentages) of cis and trans biweight mid-correlations in each correlation band, comparing all ARIES and BiB datasets.

| Correlation band | ARIES Birth |  | ARIES 7 years |  | ARIES 15-17 years |  | BiB Birth: White British |  | BiB Birth: Pakistani |  |
| --- | --- | --- | --- | --- | --- | --- | --- | --- | --- | --- |
|  | *Cis*<br>(total n=<br>119720281) | *Trans*<br>(total n=<br>77830184780) | *Cis*<br>(total n=<br>119720281) | *Trans*<br>(total n=<br>77830184780) | *Cis*<br>(total n=<br>119720281) | *Trans*<br>(total n=<br>77830184780) | *Cis*<br>(total n=<br>104564674) | *Trans*<br>(total n=<br>68269791236) | *Cis*<br>(total n=<br>104564674) | *Trans*<br>(total n=<br>68269791236) |
| -1 to -0.9 | 0<br>(0) | 3<br>(3.9e-09) | 7<br>(5.8e-06) | 0<br>(0) | 7<br>(5.8e-06) | 0<br>(0) | 0<br>(0) | 861<br>(1.3e-06) | 0<br>(0) | 298<br>(4.4e-07) |
| -0.9 to -0.8 | 3<br>(2.5e-06) | 401<br>(5.2e-07) | 35<br>(2.9e-05) | 359<br>(4.6e-07) | 35<br>(2.9e-05) | 177<br>(2.3e-07) | 45<br>(4.3e-05) | 24088<br>(3.5e-05) | 16<br>(1.5e-05) | 9086<br>(1.3e-05) |
| -0.8 to -0.7 | 43<br>(3.6e-05) | 4470<br>(5.7e-06) | 116<br>(9.7e-05) | 39097<br>(5.0e-05) | 60<br>(5.0e-05) | 1075<br>(1.4e-06) | 403<br>(3.9e-04) | 175848<br>(2.6e-04) | 196<br>(1.9e-04) | 80979<br>(1.2e-04) |
| -0.7 to -0.6 | 979<br>(8.2e-04) | 922237<br>(0.001) | 4742<br>(0.004) | 4795746<br>(0.006) | 322<br>(2.7e-04) | 183597<br>(2.4e-04) | 3261<br>(0.003) | 2729487<br>(0.004) | 1325<br>(0.001) | 881459<br>(0.001) |
| -0.6 to -0.5 | 36468<br>(0.031) | 42974963<br>(0.06) | 81067<br>(0.07) | 95777908<br>(0.1) | 15467<br>(0.01) | 19415165<br>(0.02) | 48650<br>(0.05) | 53148470<br>(0.08) | 24738<br>(0.02) | 26749688<br>(0.04) |
| -0.5 to -0.4 | 274527<br>(0.23) | 317273367<br>(0.4) | 404340<br>(0.3) | 450807392<br>(0.6) | 173905<br>(0.1) | 206058410<br>(0.3) | 267285<br>(0.3) | 282045322<br>(0.4) | 191497<br>(0.2) | 208213229<br>(0.3) |
| -0.4 to -0.3 | 959877<br>(0.8) | 1001068181<br>(1.3) | 1171964<br>(1) | 1150801019<br>(1.5) | 748813<br>(0.6) | 753206253<br>(1) | 867102<br>(0.8) | 828046988<br>(1.2) | 742056<br>(0.7) | 729079177<br>(1.1) |
| -0.3 to -0.2 | 2879403<br>(2.4) | 2593874102<br>(3.3) | 3345991<br>(2.8) | 2880767119<br>(3.7) | 2628050<br>(2.2) | 2242288139<br>(2.9) | 2673802<br>(2.6) | 2270361950<br>(3.3) | 2468830<br>(2.4) | 2129015877<br>(3.1) |
| -0.2 to -0.1 | 10426258<br>(8.7) | 8147044273<br>(10.5) | 11178176<br>(9.3) | 8693771577<br>(11.2) | 10545834<br>(8.8) | 8106743185<br>(10.4) | 9759864<br>(9.3) | 7355856656<br>(10.8) | 9444571<br>(9) | 7138202626<br>(10.5) |
| -0.1 to 0 | 30864827<br>(25.8) | 21476134284<br>(27.6) | 30021734<br>(25.1) | 20523054551<br>(26.4) | 30539452<br>(25.5) | 21542953870<br>(27.7) | 30546614<br>(29.2) | 20177889631<br>(29.6) | 31356436<br>(30) | 20543387044<br>(30.1) |
| 0 to 0.1 | 40717688<br>(34) | 23557573193<br>(30.3) | 39800719<br>(33.2) | 22864354192<br>(29.4) | 42748394<br>(35.7) | 24869018033<br>(32) | 36475305<br>(34.9) | 21656131682<br>(31.7) | 37728258<br>(36.1) | 22534057813<br>(33) |
| 0.1 to 0.2 | 22386616<br>(18.7) | 12933163606<br>(16.6) | 21062427<br>(17.6) | 12473794995<br>(16) | 21802894<br>(18.2) | 12999434519<br>(16.7) | 14832604<br>(14.2) | 9391333794<br>(13.8) | 14258103<br>(13.6) | 9159570056<br>(13.4) |
| 0.2 to 0.3 | 6863876<br>(5.7) | 4596105049<br>(5.9) | 7358153<br>(6.2) | 4892148390<br>(6.3) | 6555957<br>(5.5) | 4359633908<br>(5.6) | 4831645<br>(4.6) | 3394420353<br>(5) | 4447992<br>(4.3) | 3163558256<br>(4.6) |
| 0.3 to 0.4 | 2564609<br>(2.1) | 1875571273<br>(2.4) | 2997820<br>(2.5) | 2120261423<br>(2.7) | 2447776<br>(2) | 1716228979<br>(2.2) | 2188682<br>(2.1) | 1521328435<br>(2.2) | 1978938<br>(1.9) | 1396655448<br>(2.1) |
| 0.4 to 0.5 | 1070912<br>(0.9) | 821068527<br>(1.1) | 1349404<br>(1.1) | 1011296118<br>(1.3) | 969174<br>(0.8) | 685508087<br>(0.9) | 1074701<br>(1) | 722291874<br>(1.1) | 982620<br>(0.9) | 665865317<br>(1) |
| 0.5 to 0.6 | 460221<br>(0.38) | 336608066<br>(0.4) | 626390<br>(0.5) | 467873540<br>(0.6) | 371346<br>(0.3) | 241537376<br>(0.3) | 542477<br>(0.5) | 349881373<br>(0.5) | 506367<br>(0.5) | 322746560<br>(0.5) |
| 0.6 to 0.7 | 167909<br>(0.14) | 106864972<br>(0.1) | 247738<br>(0.2) | 165454442<br>(0.2) | 131295<br>(0.1) | 70360685<br>(0.09) | 296157<br>(0.3) | 179320379<br>(0.3) | 278815<br>(0.3) | 167808304<br>(0.2) |
| 0.7 to 0.8 | 41995<br>(0.04) | 22537164<br>(0.03) | 62118<br>(0.05) | 32873193<br>(0.04) | 37271<br>(0.03) | 16648866<br>(0.02) | 135469<br>(0.1) | 75362451<br>(0.1) | 131674<br>(0.1) | 73361627<br>(0.1) |
| 0.8 to 0.9 | 3832<br>(0.003) | 1396572<br>(0.002) | 6816<br>(0.006) | 2312808<br>(0.003) | 3814<br>(0.003) | 964425<br>(0.001) | 20326<br>(0.02) | 9438017<br>(0.01) | 21975<br>(0.02) | 10543441<br>(0.02) |
| 0.9 to 1 | 238<br>(2.0e-04) | 77<br>(9.9e-08) | 524<br>(4.4e-04) | 911<br>(1.2e-06) | 415<br>(3.5e-04) | 31<br>(4.0e-08) | 282<br>(2.7e-04) | 3577<br>(5.2e-06) | 267<br>(2.6e-04) | 4951<br>(7.3e-06) |

### *Cis* co-methylation structure across the genome (as measured by the 450k array)

To investigate the influence of physical distance on the *cis* co-methylation structure in more detail, decay plots were created across all autosomes (as measured by the 450k array) separating positive and negative correlations. In line with previous work, *cis* correlations within 10kb across the whole genome reveals a smooth decay that reduces to background correlation (about 0.125) at approximately 3kb (28, 30, 34) and the decay is identical across population cohorts and across ethnic groups (32). This confirms that it is unlikely to be driven by LD, partly because the decay is over a vastly smaller distance (see LD decay plot in (54) for comparison), and partly because the decay is identical for two different ethnic groups. Furthermore, the decay shows a high degree of heterogeneity and is identical between birth and adolescence, meaning *cis* correlation does not solely depend on physical distance (Figure 2, supplementary figure 2). In contrast to positive correlations, we also found for the first time that negative correlations are not distance dependent as immediately adjacent sites show the lowest correlations and have a slight peak between 2.5 and 4kb.

**Figure 2:**
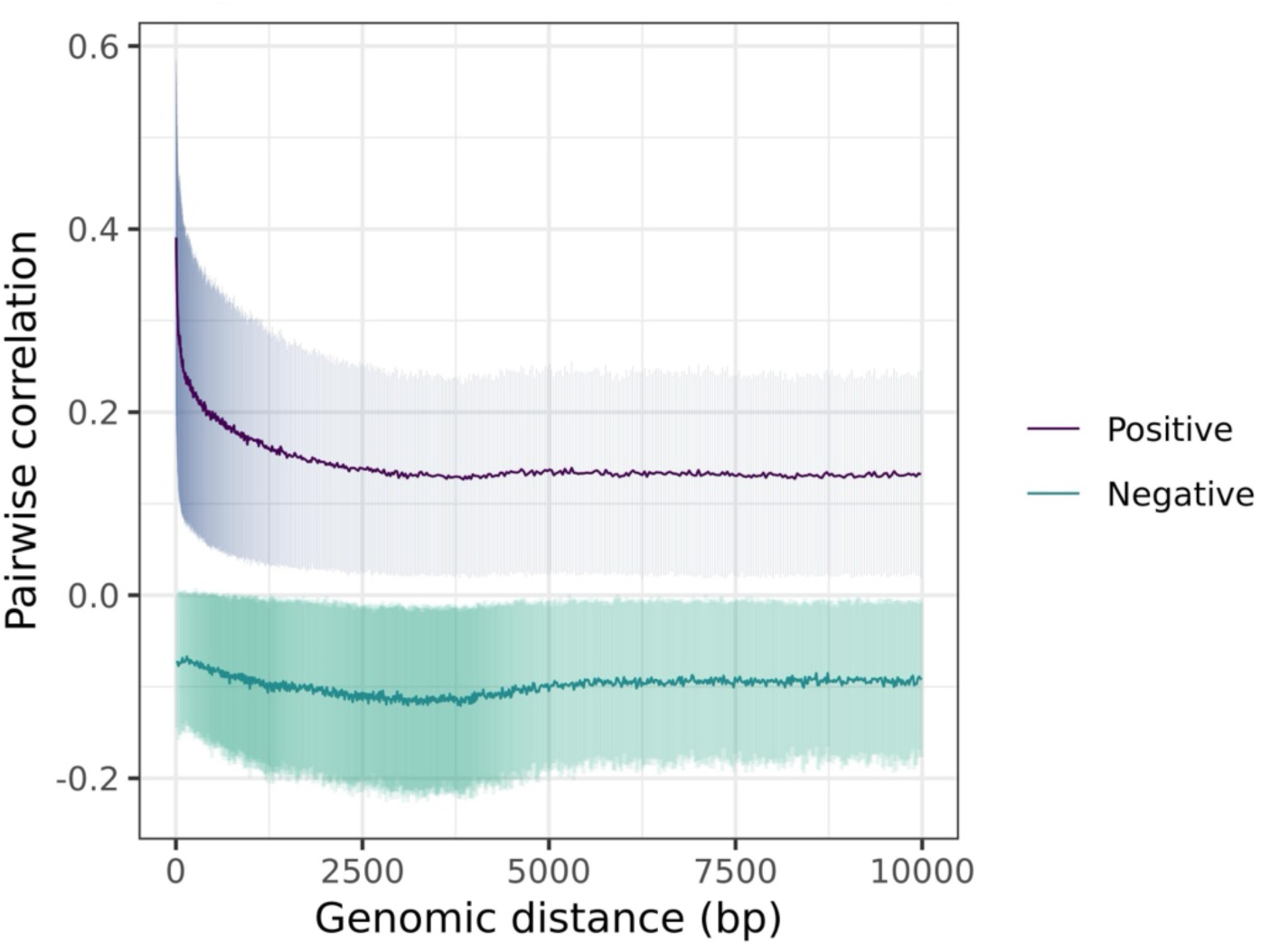
Decay plots of pairwise cis correlations (n=3,114,257, calculated using the biweight mid-correlation) from all filtered sites within 10kb of each other on the 450k array, across all autosomes (in ARIES 7 year olds; plots for all datasets can be found in supplementary figure 2). The variance in each bin (represented here by the bin standard deviation) was added to the plot to demonstrate the heterogeneity around the binned estimates.

### Stability of correlations across time, datasets, and ethnic groups

To assess whether strong correlations between DNAm sites (R>0.8, n=2,483,055 to 13,145,092 across the 5 datasets) differ across time, cohorts, and ethnic groups, we plotted mean difference (i.e. for each CpG plotting mean correlation vs difference in correlation between two groups). We found the 95% confidence intervals include a mean correlation change of zero in all tests, so we do not find strong evidence of a difference in the strength of either *cis* or *trans* correlations between any of our datasets. This suggests that correlations R>0.8 between DNAm sites are relatively stable (Table 2; supplementary figure 3). There are smaller mean changes in correlation between birth and 7 years (−0.015 for *cis*; 0.006 for *trans*) than between 7 years and adolescence (0.023 for *cis*; 0.028 for *trans*), but there is insufficient evidence to suggest that co-methylation changes with age.

**Table 2:** Mean differences in strong correlations between timepoints and datasets (defined as r>0.8, n of correlations = 2,483,055 to 13,145,092), and 95% confidence intervals (CI), for cis and trans correlations.

|  | Mean difference (*cis*) | 95% CI (*cis*) | Mean difference (*trans*) | 95% CI (*trans*) |
| --- | --- | --- | --- | --- |
| ARIES birth [reference] vs BiB birth (white British) | 0.085 | -0.05 to 0.22 | 0.088 | -0.03 to 0.2 |
| BiB birth (white British [reference] vs Pakistani) | 0.003 | -0.06 to 0.07 | 0.005 | -0.05 to 0.06 |
| ARIES birth [reference] vs 7 years | -0.015 | -0.11 to 0.08 | 0.006 | -0.04 to 0.06 |
| ARIES 7 years [reference] vs 15-17 years | 0.02 | -0.03 to 0.07 | 0.03 | -0.02 to 0.08 |

Correlations are consistently stronger in the BiB white British individuals compared to ARIES at birth (0.085 for *cis*; 0.088 for *trans*), which is reflected in the higher proportion of correlations R>0.5 in BiB identified above. In contrast, the mean difference between BiB white British and Pakistani groups is very small (0.003 for *cis*; 0.005 for *trans*), suggesting that between-cohort differences are stronger than between ethnic groups. Supplementary figure 3 shows that there are a small number of DNAm sites at which there are large changes in *trans* correlation between birth and 7 years in ARIES, between ARIES and BiB white British individuals, and between the two ethnic groups in BiB.

### Genetic and environmental influences on co-methylation

#### Influence of genetic factors on co-methylated sites

To assess whether co-methylated sites are influenced by genetic factors, we used twin heritabilities of DNAm sites (1), and we assessed the proportion of correlations which had zero, one, or two of the DNAm sites in each correlating pair associated with an mQTL (see Methods). Highly heritable DNAm sites tend to be strongly co-methylated (r>0.9), in *cis* but not *trans,* and this is consistent across datasets, with a mean heritability of 77% for DNAm sites correlated r>0.9 in *cis*. Looking at this in more detail, the heritability of both sites in highly correlating pairs (r>0.9) tend to be matched only for *cis*-correlating pairs, suggesting that very strong co-methylation is related to heritability in *cis* but not in *trans* (see Figure 3B). This may be related to genetic variants (55), as at least 84% of *cis* correlations >0.9 have both DNAm sites associated with a *cis* mQTL (not necessarily the same mQTL) across the 5 datasets (see Figure 3 and supplementary figure 4).

**Figure 3:**
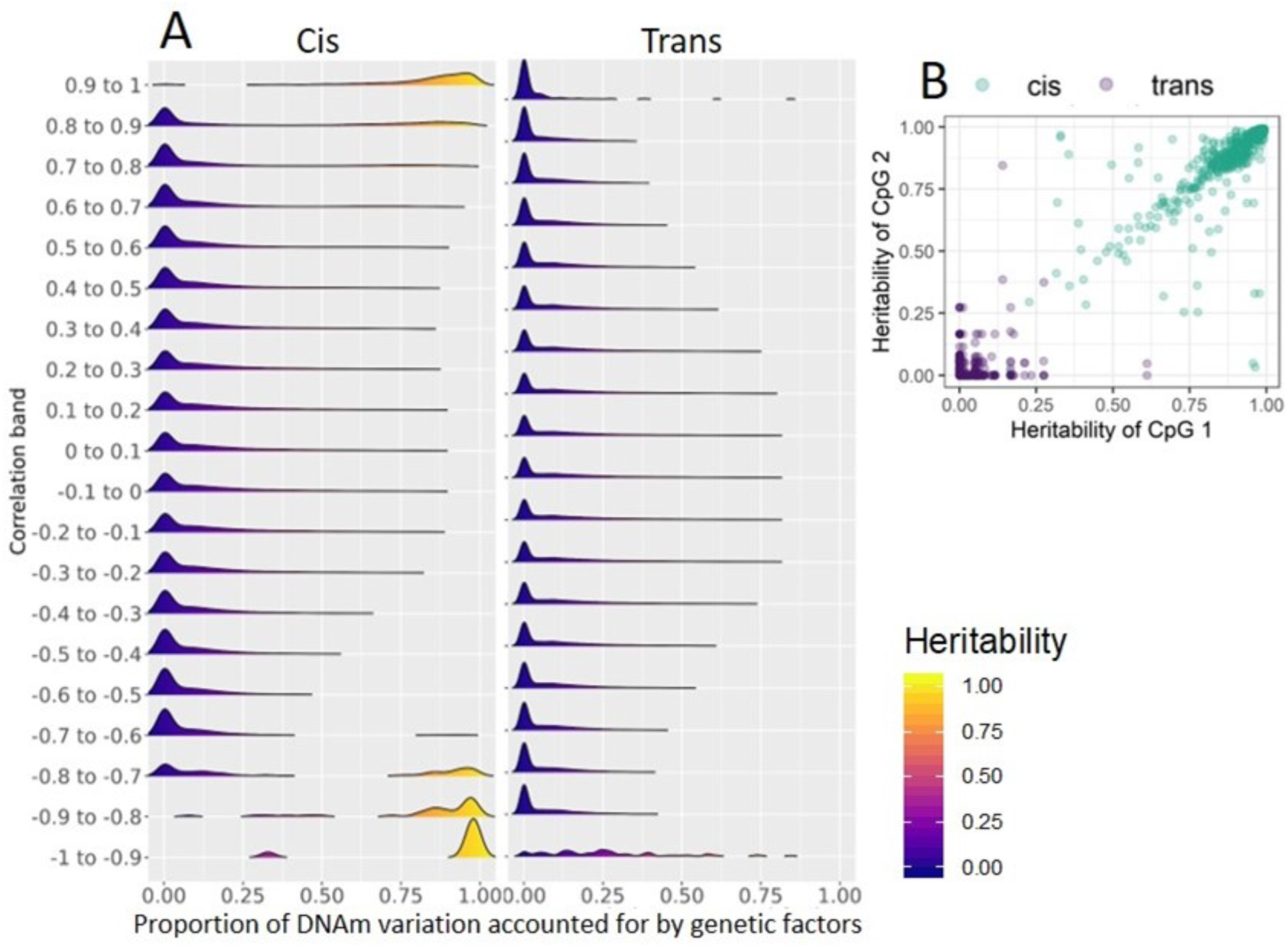
**A**: Density plots illustrating the proportion of DNAm variation due to heritability for DNAm sites with correlations of differing strengths, in ARIES 7 year olds (please see supplementary figure 4 for plots of each dataset). **B**: Scatter plot of heritability of probe pairs correlating >0.9 in ARIES 7 year olds.

To identify whether co-methylation is influenced by mQTL, we firstly assessed the proportion of correlations which had zero, one, or two of the DNAm sites in each correlating pair associated with an mQTL. At least 84% of *cis* correlations >0.9 have both DNAm sites associated with a *cis* mQTL across the 5 datasets, in line with the heritability findings above. *Trans* correlations >0.9 are most likely to have neither DNAm site associated with a *cis* mQTL in 4 of the datasets (48-55%; in the BiB white British group only 31% are associated with 0 mQTLs). This suggests that strong *trans* correlations are between sites that are less likely to have been associated with an mQTL (although it is possible they have shared genetic aetiology we have not detected - *trans* sites are less heritable and there are fewer detected *trans* mQTLs at present (56, 57)). There are slight increases in the proportion of *cis* correlations that have both DNAm sites associated with an mQTL in the ARIES adolescents. This is noticeable for *cis* correlations between −0.8 and −0.6, and between 0.8 and 0.9 (see Figure 4 and supplementary figure 5).

**Figure 4:**
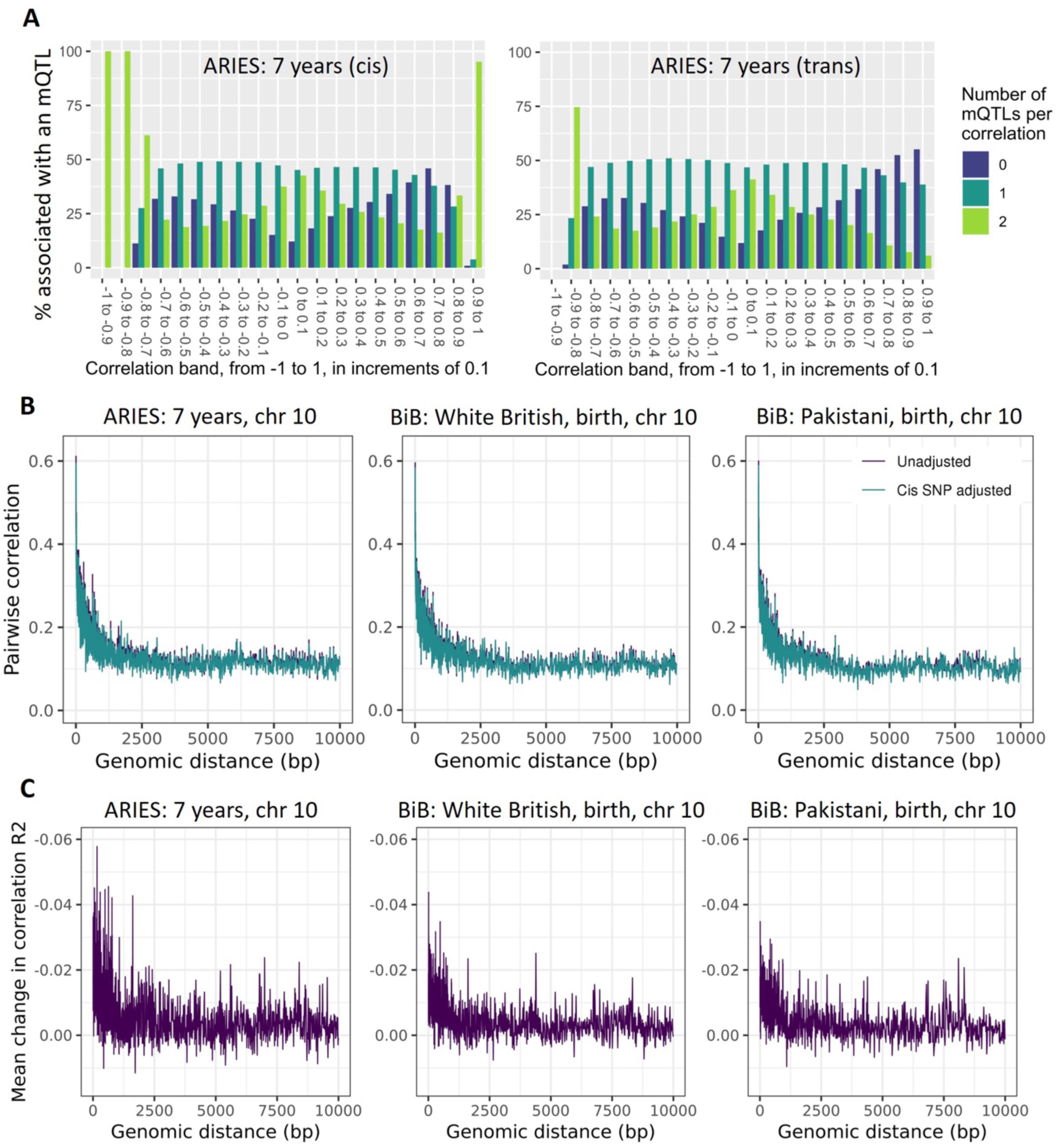
**(A)**: Bar plots of the percentage of pairwise correlations in each correlation range that have 0, 1 or 2 DNAm sites associated with a cis mQTL identified by GoDMC. Split by cis (left) and trans (right) correlating pairs, In ARIES at 7 years. **(B)**: Cis decay plot on chromosome 10 in ARIES 7 year olds, and the BiB White British and Pakistani groups (both measured in cord blood), showing the binned decay of correlation over genomic distance (purple) and the decay of correlation over genomic distance when adjusting DNAm values for the strongest associated cis SNP (green). The substantial overlap of the lines illustrates the small change in co-methylation even when adjusting for the strongest cis SNP **(C)**: Decay plot illustrating the mean change in correlation between unadjusted and strongest cis-SNP adjusted correlations.

**Figure 5:**
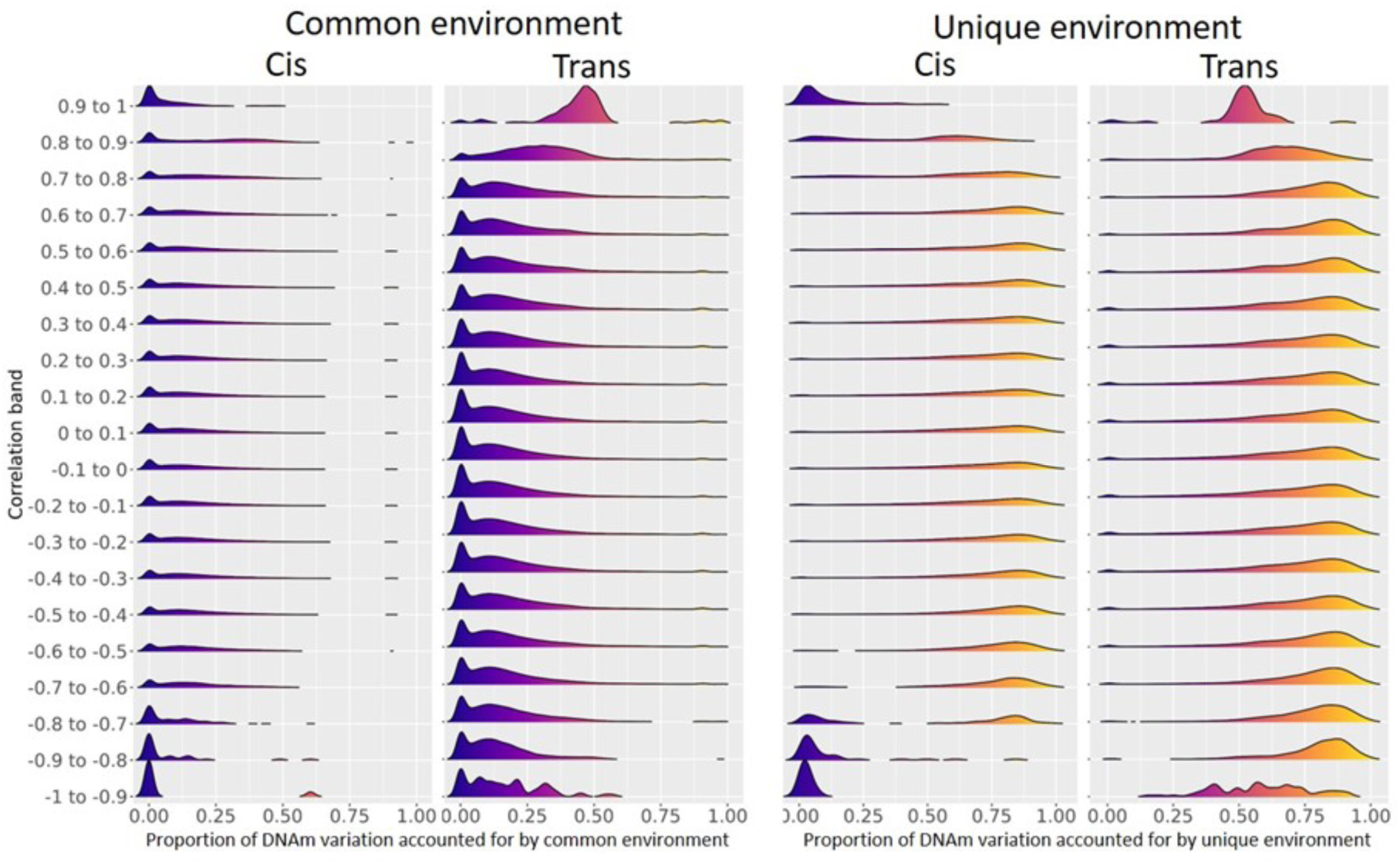
Density plots illustrating the proportion of DNAm variation due to common environment and unique environment for DNAm sites with correlations of differing strengths, in ARIES 7 year olds (please see supplementary figure 6 for plots of each dataset).

To further quantify the impact genotype may have on correlations between DNAm sites in close proximity, we assessed the extent to which *cis* SNPs impact the decay of correlation between *cis* correlating DNAm sites within 10kb. To do this we adjusted DNAm data for the strongest *cis* SNP (as identified by the largest mQTL study to date, GoDMC (56)), and reproduced the decay plot with this adjusted data (for ARIES, 11719 sites (54.8%) had an mQTL; for BiB, 11108 sites (56%)had an mQTL). We find that in both ARIES and BiB, there is limited impact of the strongest *cis* SNP on correlation structure; sites in close proximity are most affected, with a maximum reduction in bin correlation of 0.06 in ARIES 7 year olds, 0.04 in the BiB white British participants, and 0.035 in the BiB Pakistani participants. On average the correlation between sites within 1kb drops by 0.01 to 0.02, after which it plateaus to a correlation reduction of around 0.005. Of course, additive effect of multiple mQTLs may reduce correlation further than this, but here we see limited evidence of the strongest SNP affecting correlation structure. This is, to our knowledge, the first illustration of the direct impact of genetic variants on co-methylation between nearby sites. This may suggest that *cis* co-methylation also depends on environmental effects, which may act on the same pathways and in the same direction as genetic effects (58).

#### Influence of environment on co-methylated sites

Environmental influences on individual DNAm sites have been separated as common (or shared) environment and unique environment (which includes measurement error) though twin studies (1). Consistent with the high heritability estimates for sites that correlate r>0.9 in *cis*, the common and unique environment estimates for those sites are low. In *trans,* almost all sites which correlate with another DNAm site at r>0.9 are influenced on average 41% by common environment, and 49% by unique environment (which includes measurement error and interindividual stochastic variation). Because DNAm sites involved in strong co-methylation in *trans* are influenced by environmental factors, strong *trans* co-methylation may be driven by environmental influences.

### Features of highly co-methylated sites

#### Overview

Because *cis* and *trans* sites correlated >0.9 have distinct features in terms of genetic and environmental influences, we conducted an in-depth analysis to identify the biological mechanism of this co-methylation. The number of correlations R>0.9 in each dataset are detailed in Table 3. Throughout the following section for simplicity we show results for the ARIES 7 year olds, as results were very similar across all five datasets (See Supplement for other datasets).

**Table 3:** Numbers of cis and trans correlations r>0.9, and the number of unique DNAm sites these correlations are between.

| Cohort | Timepoint | *Cis* correlations | *Trans* correlations |
| --- | --- | --- | --- |
| ARIES | Birth (n=849) | 238 (305 sites) | 77 (46 sites) |
|  | 7 years (n=910) | 524 (589 sites) | 911 (162 sites) |
|  | 15-17 years (n=921) | 415 (486 sites) | 31 (24 sites) |
| BiB | Birth – white British (n=424) | 282 (413 sites) | 3577 (683 sites) |
|  | Birth - Pakistani (n=439) | 267 (387 sites) | 4951 (669 sites) |

#### Biological mechanisms of highly co-methylated sites

*Cis* and *trans* sites that have correlations >0.9 display very different patterns of variance. *Cis* sites are more variable in both mean methylation level and variance, although there is a tendency for hypo- and hyper-methylated sites to have smaller variance. *Trans* correlated sites are typically hypomethylated, and have low variance. As one might expect DNAm sites under *trans*-acting influences such as transcription factors to be hypomethylated (11, 59), this would fit the behaviour of the highly *trans*-correlated sites seen here (see Figure 6 for illustration).

**Figure 6:**
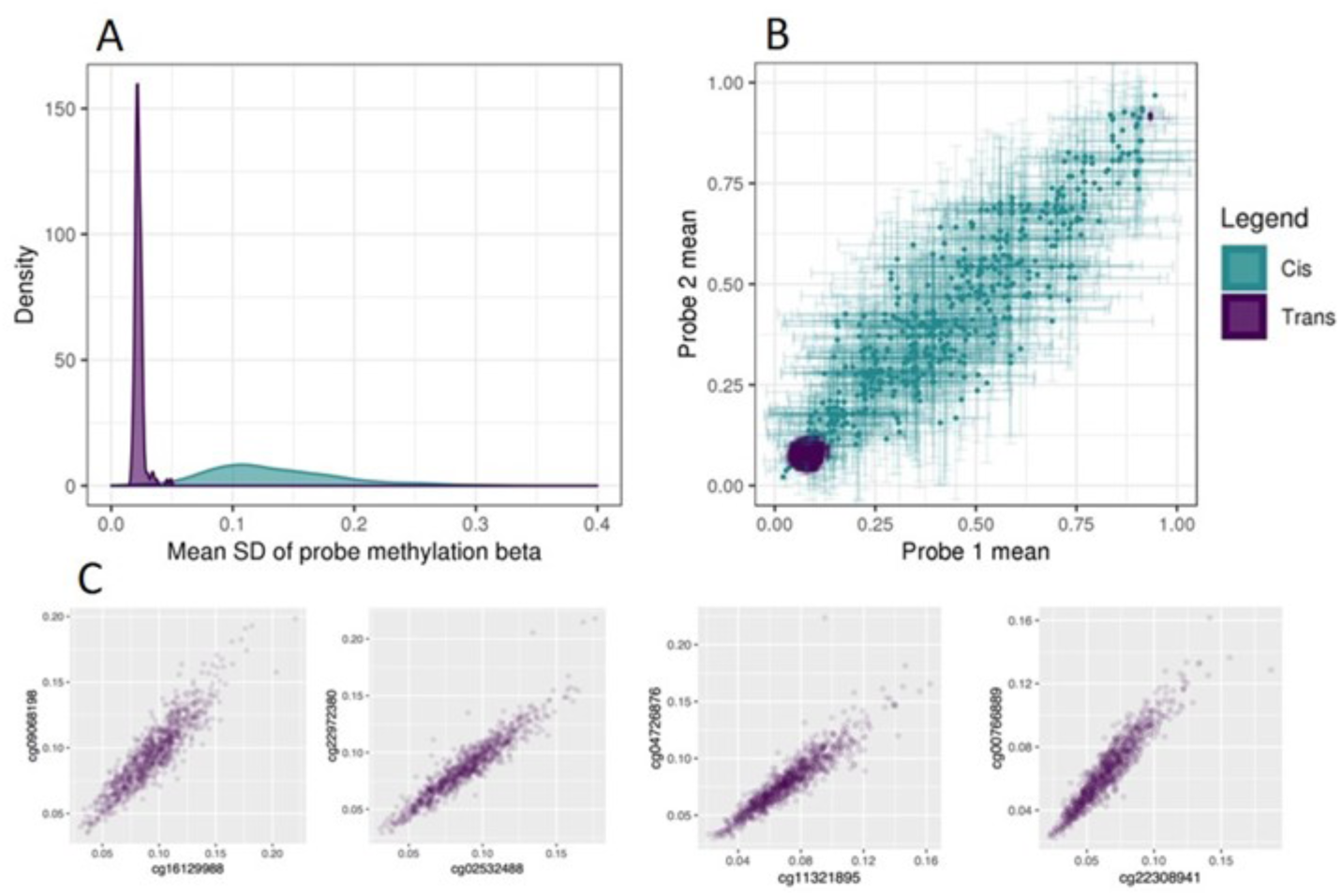
**A**: Density plot showing the mean standard deviation of cis and trans DNAm sites with correlations R>0.9. **B**: Scatter plot with error bars (standard deviation) illustrating mean methylation level of cis and trans probe pairs that correlate R>0.9. **C**: Illustration of raw scatter plots of trans-correlating probe pairs R>0.9.

#### *Trans* co-methylation structure

The architecture of co-methylation between sites that are either distant or on different chromosomes has not previously been well characterised. We find that in all datasets these sites are distributed across the genome, and are interconnected (illustrated by the circos and network plots in Figure 7 and supplementary figures 8 and 9). There is a much higher number of correlations r>0.9 in BiB than in ARIES (as shown in Table 1 and the section on overall co-methylation structure). This can be explained by correlations in BiB being higher than in ARIES across the whole distribution (see Table 1 and Table 2); something that may be due to array or sample type effects, or phenotypic plasticity. The connections between the sites correlating r>0.9 resemble scale-free networks (60, 61) with a small number of ‘hub’ nodes having large numbers of connections. All datasets except ARIES 15-17 years have a power-law alpha between 2 and 3, which is indicative of a scale-free network (60, 62) (however at 15-17 years the number of nodes is too small to assess scale-free properties appropriately). The degree distribution plots are found in supplementary figure 7.

**Figure 7:**
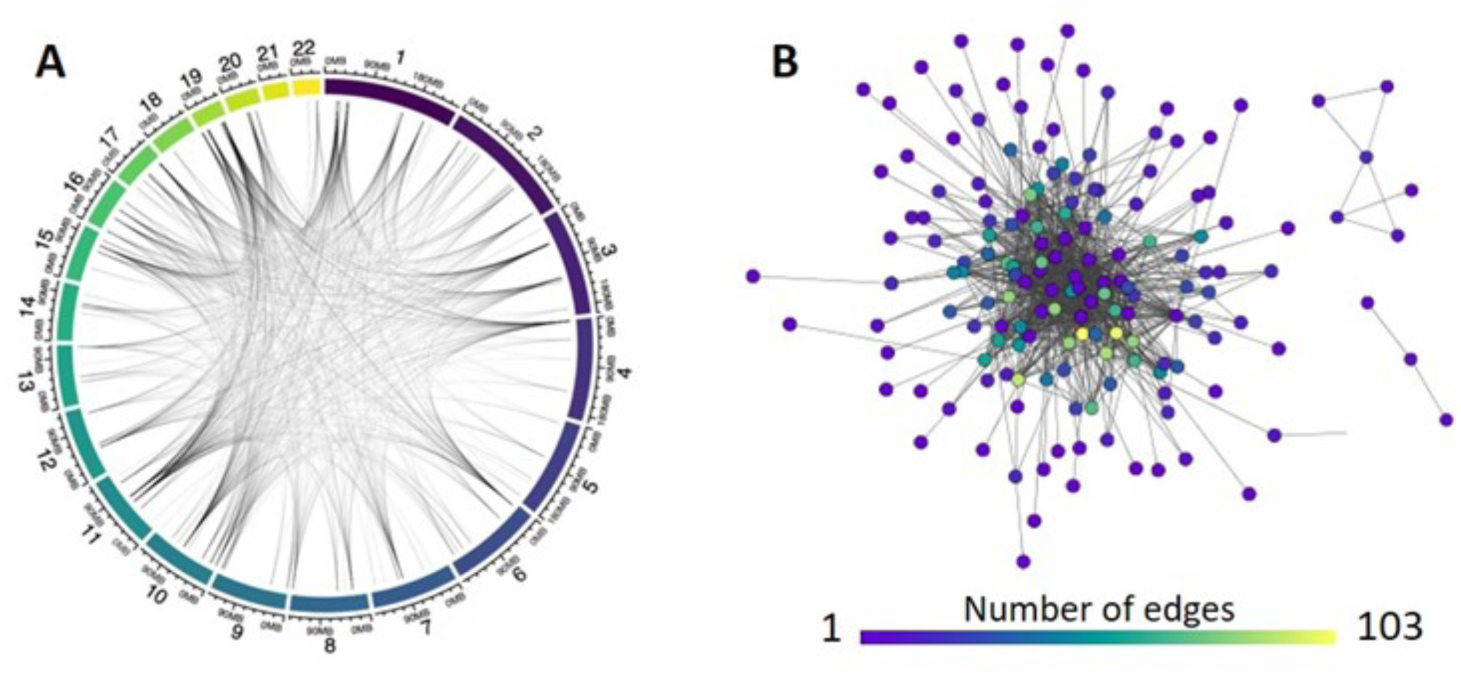
**A**: circos plot illustrating genomic distribution of trans correlations r>0.9 in ARIES 7 year olds (circos plots for all datasets can be found in supplementary figure 8). **B**: cytoscape network plot illustrating network connectivity of trans correlations r>0.9 in ARIES 7 year olds (cytoscape network plots for all datasets can be found in supplementary figure 9).

To test the preservation of network architecture for DNAm sites correlating in *trans* >0.9, we tested preservation of both the DNAm sites (nodes) and the connections between them (edges) between the five datasets. To do this we utilised the Cytoscape Network Analyzer to test the intersection between pairs of networks. Using the Binomial test we find strong evidence of preservation of both the nodes and the edges of the networks for all four comparisons; between ARIES and BiB white British cord blood, between BiB white British and Pakistani cord blood, between birth and 7 years in ARIES, and between 7 years and 15-17 years in ARIES. Results are detailed in Table 4.

**Table 4:** Node and edge intersections for trans-correlating DNAm sites r>0.9. Nodes are DNAm sites and edges are the specific pairwise correlations between two nodes. Intersection means the node, or the specific correlation between two nodes, is present in both datasets A and B. N = number of intersections; p-values assess whether there are more overlaps than expected by chance, using the Binomial exact test.

| Dataset A | Dataset B | Dataset A nodes (edges) $R > 0.9$ | Dataset B nodes (edges) $R > 0.9$ | Intersecting nodes N (p-value) | Intersecting edges N (p-value) |
| --- | --- | --- | --- | --- | --- |
| ARIES at birth | BiB white British at birth | 46 (77) | 683 (3577) | 31 (7.7e-39) | 9 (1.3e-04) |
| BiB white British at birth | BiB Pakistani at birth | 683 (3577) | 669 (4951) | 401 (<2.2e-308) | 1056 (<2.2e-308) |
| ARIES at birth | ARIES 7 year olds | 46 (77) | 162 (911) | 35 (8.6e-47) | 23 (9.3e-19) |
| ARIES 7 year olds | ARIES 15-17 year olds | 162 (911) | 24 (31) | 21 (4.3e-31) | 15 (1.9e-16) |

#### Enrichments of *cis* co-methylated sites

To illustrate the potential utility of strong *cis* co-methylation, enrichment analyses were conducted to assess the enrichment for DNAm sites being located within specific transcription factor binding sites (TFBS), chromatin states, and genomic regions. All pairs of DNAm sites correlating R>0.9 and within 1Mb of each other were included in this analysis. Enrichments were virtually identical for all 5 datasets. *Cis* correlating sites are strongly enriched for chromatin states associated with poised promoters (PromP; OR=3.9 to 4.8, p=1.1e-09 to 2.4e-17), and less strongly enriched for weak enhancers (EnhW1; OR=2.1 to 2.7, p=1.4e-03 to 8.4e-05, and EnhW2; OR=1.9 to 2.2, p=0.03 to 0.02). *Cis* correlating DNAm sites are strongly enriched for a select few TFBS in blood: RNA Polymerase III (Pol3) (OR=10.3 to 18.7, p=2.4e-05 to 4.6e-09), BRF1 (OR=10.5 to 18.5, p=2.9e-05 to 5e-09), and BDP1 (OR=6.8 to 11.4, p=2e-04 to 1.5e-06), with weaker enrichment of TFIIIC-110 and RPC155. Heatmaps of TFBS and chromatin enrichments are shown in Figure 8. *Cis* sites showed strongest enrichment for location in promoters (OR=1.62 to 2.5, p=6.2e-06 to 1.1e-17) and 5’UTR (OR=1.6 to 2, p=0.02 to 9.8e-09), with weaker enrichment in exons (OR=1.4 to 1.6, p=0.02 to 9.7e-06) (shown in supplementary figure 10). As BRF1 and BDP1 are essential for RNA polymerase III transcription, these enrichments suggest that coordination of methylation state of sites in close proximity is primarily a feature of active promoter regions involved in regulation of short RNAs essential for cellular function transcribed by RNA polymerase III.

**Figure 8:**
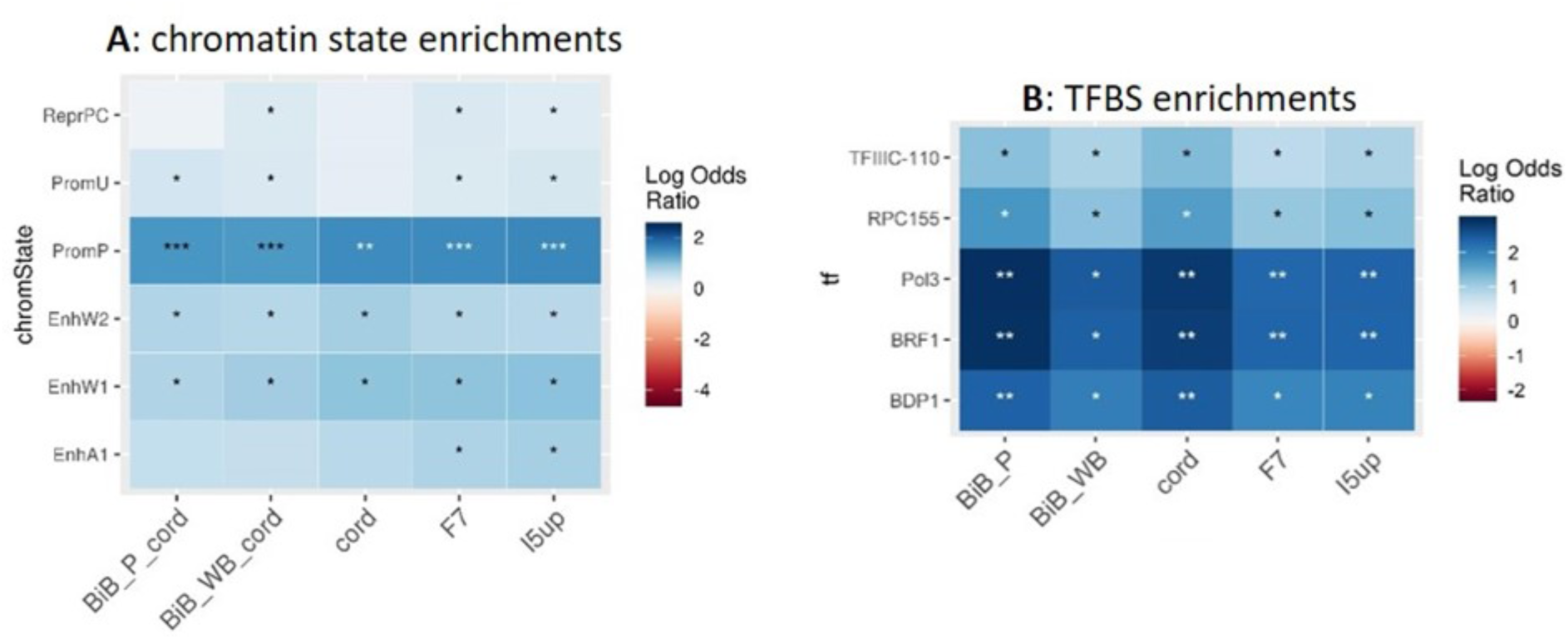
Enrichments of cis-correlating sites r>0.9 for **A** chromatin states and **B** transcription factor binding sites (TFBS). Enrichment analyses were conducted using LOLA, with enrichments in blood only.; *p<0.05 **p<1e-05 ***p<1e-10; asterisk colour differences are to help visibility.

**Figure 9:**
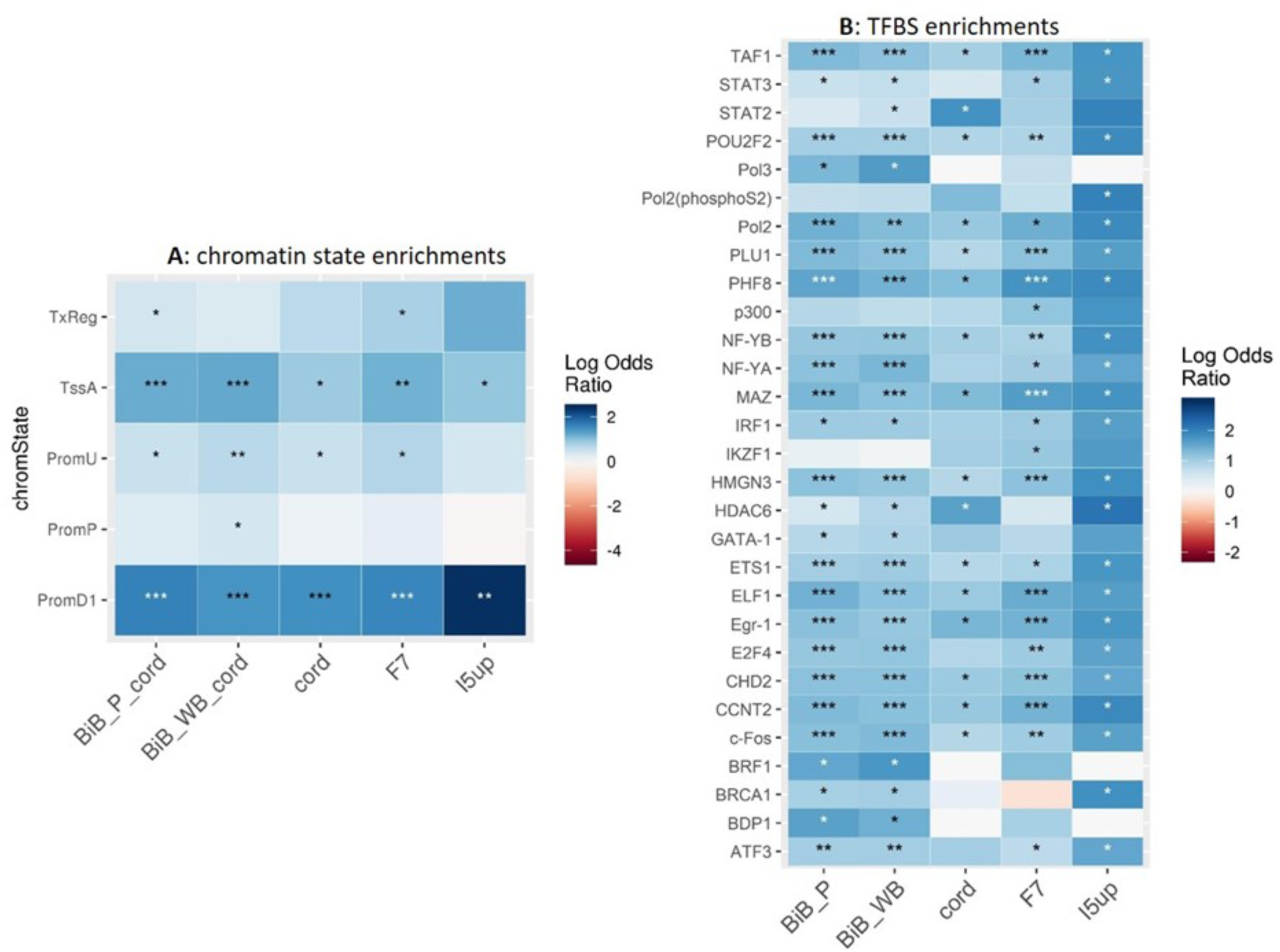
Enrichments of trans-correlating sites r>0.9 for **A** chromatin states and **B** transcription factor binding sites (TFBS). Enrichment analyses were conducted using LOLA, with enrichments in blood only.; *p<0.05 **p<1e-05 ***p<1e-10

#### Enrichments of *trans* co-methylated sites

Next, we tested whether sites involved in strong (R>0.9) *trans* correlations were enriched for locations in TFBS, chromatin states and genomic regions. *Trans* correlating sites are enriched for 29 of the tested TFBS: including Pol2 (OR=3.1 to 6.5, p=2.9e-03 to 5.2e-12), PHF8 (OR=3.6 to 6.4,p=3.2e-05 to 9.9e-81), MAZ (OR=3.3 to 5.7,p=9.3e-05 to 8.9e-63), ELF1 (OR=3 to 5,p=5.1e-04 to 3.6e-63), Egr-1 (OR=3.1 to 5.6, p=3e-04 to 2.5e-49), and TAF1 (OR=2.7 to 5.6, p=1.8e-03 to 9.5e-47). In contrast to *trans*-mQTL associated sites (56), we do not see enrichment for cohesin related TFs, which may reflect the small proportion of *trans* correlations found on the same chromosome. *Trans*-correlating sites are strongly enriched for chromatin states associated with promotors downstream of transcription start sites 1 (PromD1) (OR=4.1 to 11, p=5.1e-07 to 1.9e-78), and active transcription start sites (TssA) (OR=2.6 to 3.4, p=4.9e-02 to 1e-30), and weakly enriched for locations at promotors upstream of transcription start sites (PromU) (OR=1.6 to 2, p=0.26 to 1.8e-09). *Trans* sites show no evidence of enrichments in the adolescents in ARIES; for all other datasets we see the strongest enrichment for CpG islands (OR=2.2 to 6.1, p=3.8e-08 to 5.7e-28), with moderate enrichment in promoters (OR=1.9 to 2.6, p=4e-04 to 1.3e-32) and 5’UTR (OR=2.1 to 2.2, p=3.3e-05 to 4.7e-21) (shown in supplementary figure 11). As such, co-methylation of *trans* sites may relate to active transcription.

#### Enrichment of inter-chromosomal chromatin contacts in *trans* co-methylated sites

To identify whether strong *trans* correlations are located in sites where chromatin contacts are formed, we assessed the overlap of the *trans* correlations r>0.9 with the (Rao et al., 2014) Hi-C data. This analysis was performed with the 860 inter-chromosomal *trans* correlations r>0.9 in the ARIES 7 year olds, and the 3577 in the BiB white British dataset. We find a strong enrichment of inter-chromosomal Hi-C contacts in the real correlation data as compared to 1000 permutations of the data, with no permutation set having a higher count of overlaps than the real data in either ARIES or BiB (n=46 in ARIES and n=654 in BiB; p=<2.2e-308 for both datasets) (Figure 10). This suggests that correlation of DNAm sites across chromosomes is at least in part likely to be related to inter-chromosomal contacts, which would mean that coordinated methylation states between DNAm sites on different chromosomes could be functionally relevant to inter-chromosomal chromatin contacts.

**Figure 10:**
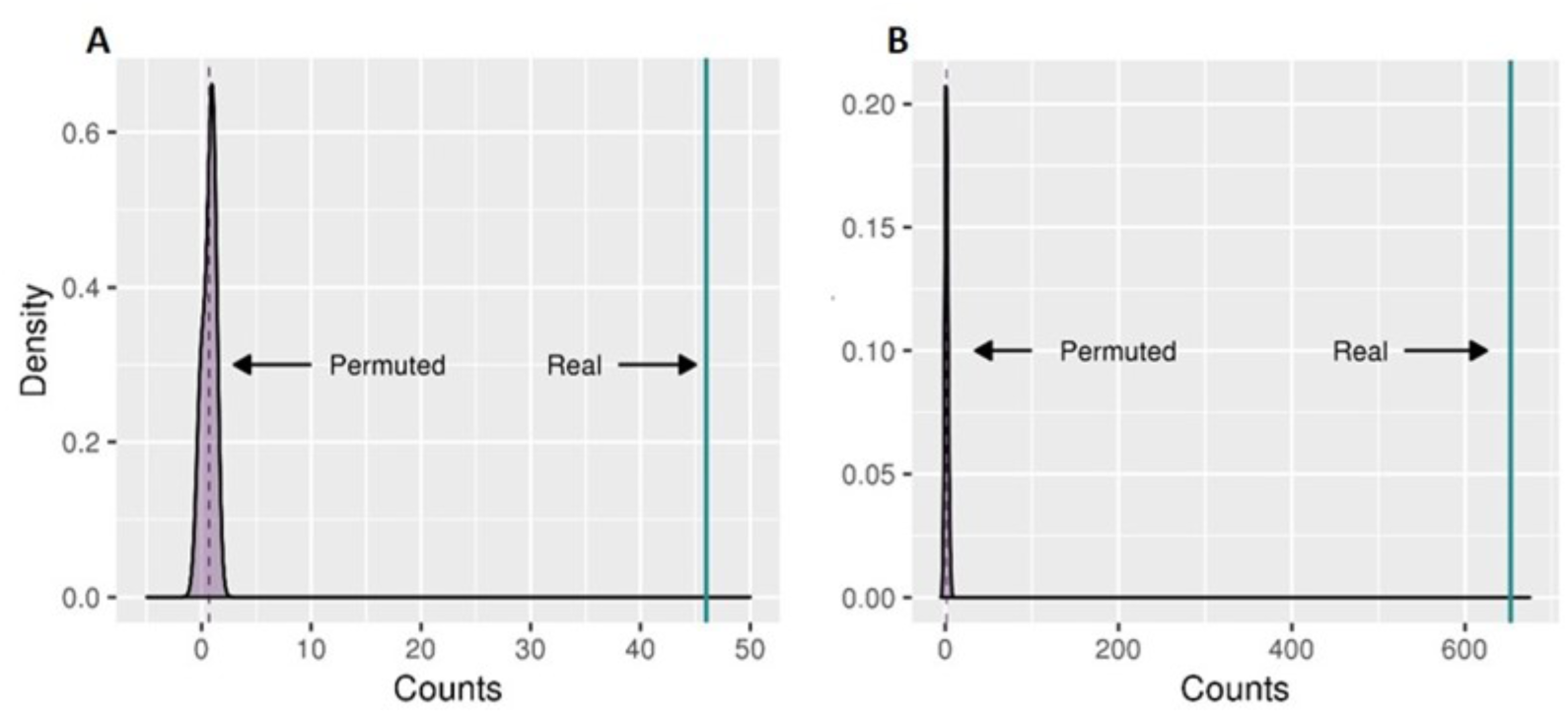
**A**: In ARIES the overlap in the permuted datasets is either 0 or 1, whereas there are 46 overlaps in the real data. **B**: In BiB the overlap in the permuted datasets is between 0 and 5, whereas there are 654 overlaps in the real data.

## Discussion

This work has demonstrated that DNAm data has a stable co-methylation structure, both in *cis* and in *trans*, that persists with aging from birth to 15-17 years, and is similar across White British people living in two different geographical areas, one which includes participants that on average are more socioeconomically advantaged than the national and local area average (63, 64), with the participants born in the early 1990s (ARIES); and the other from a more deprived area with participants born 2007-10 (BiB). In BiB there was stability between White British and Pakistani subgroups. It suggests that co-methylation of DNAm sites may be related to specific aspects of genome regulation, where sites in *cis* are fundamentally different to co-methylation of sites which are distant or on different chromosomes. This includes the novel observation that *cis* co-methylation structure is not generally substantially reduced when adjusting for the strongest *cis*-mQTL.

Through enrichment analyses we find that co-methylation is likely to be related to short RNA transcription associated with RNA polymerase III in *cis*; this might indicate that co-methylation is driven by environmental effects or TFs, which could be correlated with genetic effects (58). We have demonstrated in humans that *trans*-correlating DNAm sites are likely to represent inter-chromosomal contacts or regulation by multiple transcription factors, and thus they are likely to represent shared regulation. This is consistent with recent published evidence of correlation between DNAm sites in inter- and intra-chromosomal chromatin contact regions in mice (16).

This work suggests that DNA co-methylation is relatively stable across the groups that we have included in this study. In all groups, co-methylation is weak (R <0.2,>-0.2) between the vast majority of DNAm sites (83-87%), with a greater proportion of *cis* than *trans* co-methylated sites having very strong (>0.9) co-methylation, illustrating the role of close physical proximity in co-methylation. To test how well co-methylation between specific sites replicated, we restricted to correlations >0.8 in each dataset, and found that the magnitudes of correlation for sites in *cis* and *trans* are similar between ARIES and BiB White British (cord blood), and between White British and Pakistani (cord blood) in BiB. *Cis* and *trans* correlations were also similar in ARIES between birth and age 7 (with *cis* and *trans* mean differences close to the null), and between age 7 and 15 there appears to be a slight increase in correlation. However, we acknowledge that some of these differences in means had wide confidence intervals. The stability we have identified is important because it means these DNAm sites might also be reliably co-methylated in other datasets - this would mean co-methylation structure could have the potential to be more broadly applied to EWAS and DMR analyses. However this would need to be tested in older age groups, in more social groups including a greater range of ethnic groups, and in individuals born and residing outside of the UK.

We demonstrate that although *cis* co-methylation is distance-based, there is a large degree of variation. This is likely to reflect the fact that we find strong *cis* correlations are related to genomic regions (most strongly promotors and 5’UTRs). Although we show that DNAm sites that have strong *cis* correlations are highly heritable and are associated with *cis* mQTLs, we have shown that adjusting for the strongest *cis* mQTL does not substantially impact *cis* co-methylation structure; this is consistent with co-methylation structure not mirroring LD. Although we show there are differences in co-methylation between the five datasets we use in this study, we found that the biological meaning behind *cis* co-methylation structure is consistent across studies. From our enrichment analysis it appears that *cis* co-methylation is enriched for being located only at binding sites of transcription factors essential for RNA polymerase III transcription; this is supported by the recent demonstration that *cis* mQTL SNPs overlap TFBS (7, 56). Coordinated DNAm states of locations in close proximity are therefore likely to be involved in regulation of the transcription of short RNAs involved in essential cellular functions (such as protein synthesis and transport) (65, 66), and, most relevant to the datasets we use in this study, RNA polymerase III transcription has been shown to be a determinant of growth of both the cell and the organism (67).

We also found consistent biological enrichment between datasets for DNAm sites at least 1Mb apart that are strongly co-methylated. Strong correlations between DNAm sites on different chromosomes are shown here to be strongly enriched for inter-chromosomal contacts. Coordinated methylation states have been shown previously between contacting inter-chromosomal regions of the genome in mice (16); we demonstrate that this phenomenon can be identified using existing DNAm microarray and Hi-C data in humans. This is very much in line with recent work which shows co-methylated DNAm sites on the same chromosome are enriched for chromosomal loop contact sites (36). One might in fact expect the true number of inter-chromosomal contacts between highly co-methylated inter-chromosomal DNAm sites to be higher than reported here, as methods such as Hi-C do not pick up many inter-chromosomal contacts due to the greater distance between them than between *cis* contact sites (68). The enrichment for a multitude of TFBS suggests inter-chromosomal co-methylation may be related to transcription factor networks, which have key roles in genome regulation (69) and have been shown to mediate inter-chromosomal chromatin contacts (70). Finally, highly co-methylated *trans* sites are influenced almost entirely by non-genetic factors – that makes co-methylation an interesting factor to study in relation to environmental influences on genome regulation, given the chromatin and transcription factor enrichments for these sites.

We acknowledge that there are some limitations to this work. The first is that our datasets only span birth to adolescence; we do not know how applicable our results will be to adults. Differences between cohorts (ARIES and BiB cord blood datasets) are likely to have been influenced by the use of different array platforms, smaller numbers of BiB participants, and diversity of sample types in ARIES and phenotypic plasticity which may resulted in more correlations >0.5 in both groups of BiB than in ARIES. This study only considered DNAm in blood - DNAm is cell-type- and tissue-specific, and as such the applicability of these results to DNAm in other tissues may be fairly limited (71); although it is likely to be better for the highly heritable strong *cis* correlations (1). Of particular importance is a demonstration that *trans*-chromosomal promoter contacts have been shown to be cell-type specific (72), and so future work may benefit from utilisation of less heterogeneous cell type populations than blood or advanced deconvolution methods to investigate the functions of trans-chromosomal co-methylation. Finally, our study utilised DNA methylation arrays, which measure only 2-4% of DNAm sites and contain an over-representation of specific types of genomic regions, including CpG island regions, promotors, and enhancers. Whilst we have corrected for this overrepresentation in our downstream analyses, future work needs to ascertain whether these conclusions hold in genome-wide data.

## Conclusions

DNAm has a stable co-methylation structure in humans that persists to at least adolescence and across social groups (here represented by both cohort and ethnicity). *Cis* co-methylation is likely to be related to short RNA transcription that is associated with RNA polymerase III. *Trans* co-methylation is highly enriched in regions of inter-chromosomal contacts, and for the binding sites of multiple transcription factors, suggesting that co-methylation may have a role in 3D genome regulation.

## Materials and methods

### Participants

Data were taken from participants of two birth cohorts: the Avon Longitudinal Study of Parents and Children (ALSPAC) (63, 64), and the Born in Bradford study (BiB) (73) (Table 5). Detailed descriptions of the cohorts can be found in the Supplementary Methods (63, 64, 73);. In brief, ALSPAC is a multi-generational cohort study based in the Bristol area, comprising mostly white British participants. The original cohort were 14,541 pregnancies with a predicted delivery date between April 1991 and December 1992. A subsample of participants (known as ARIES) of 1022 mother-child pairs had DNAm data generated at five timepoints: birth, 7 years and 15-17 years in the children, and during pregnancy and 12-18 years later in the mothers. Our study utilises the three child timepoints only.

**Table 5:**
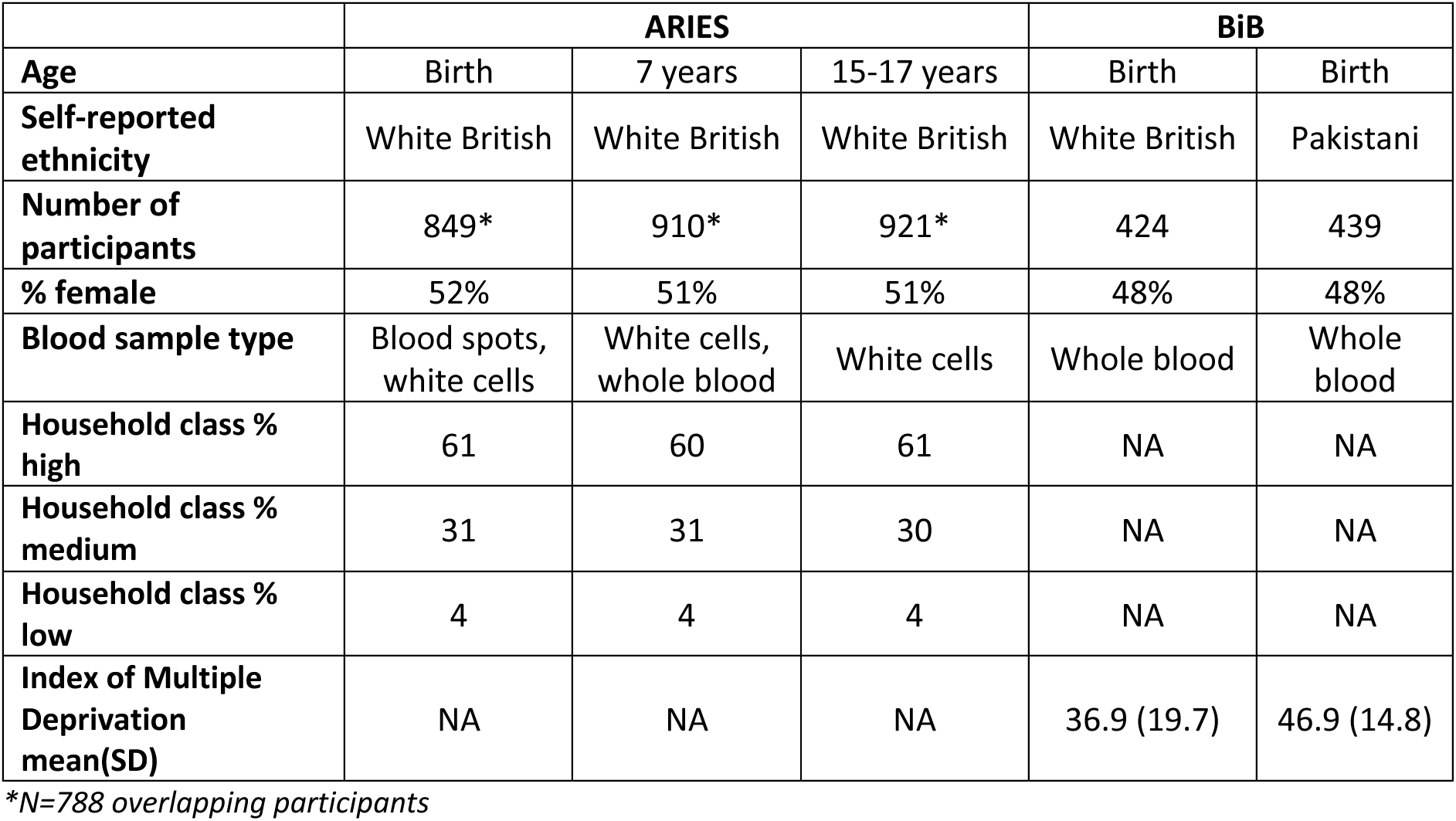
Overview of study participants and sociodemographic characteristics.

|  | **ARIES** |  |  | **BiB** |  |
| --- | --- | --- | --- | --- | --- |
| **Age** | Birth | 7 years | 15-17 years | Birth | Birth |
| **Self-reported ethnicity** | White British | White British | White British | White British | Pakistani |
| **Number of participants** | 849* | 910* | 921* | 424 | 439 |
| **% female** | 52% | 51% | 51% | 48% | 48% |
| **Blood sample type** | Blood spots, white cells | White cells, whole blood | White cells | Whole blood | Whole blood |
| **Household class % high** | 61 | 60 | 61 | NA | NA |
| **Household class % medium** | 31 | 31 | 30 | NA | NA |
| **Household class % low** | 4 | 4 | 4 | NA | NA |
| **Index of Multiple Deprivation mean(SD)** | NA | NA | NA | 36.9 (19.7) | 46.9 (14.8) |

\*N=788 overlapping participants

BiB is a longitudinal, multi-ethnic cohort study based in Bradford, UK. The original cohort were 13,776 pregnancies with a predicted delivery date between March 2007 and November 2010. Like ALSPAC it was set up to investigate factors which influence child health and development, but with a particular focus on child morbidity and mortality, as rates of these have been higher in Bradford than the rest of the UK (73). Bradford has a high rate of economic deprivation – one third of the neighbourhoods in Bradford are in the most deprived 10% of neighbourhoods in England (74, 75).

Around 20% of the population are of South Asian descent, and 90% of these individuals are of Pakistani origin (73); the BiB DNAm subsample was specifically designed to be multi-ethnic, and so of those eligible, 500 White British and 500 Pakistani mothers were selected to have DNAm generated for themselves and their children.

### DNA methylation data

DNAm data generation is described in detail for each cohort in the supplementary methods. Consent for biological samples for ARIES and BiB was collected in accordance with the Human Tissue Act (2004).

Briefly, in ARIES DNAm profiles were measured using Illumina Infinium 450k beadchip arrays (Illumina, San Diego, CA, USA). Processing, extraction, and quality control of DNAm data has been described in detail for these samples (76), as have normalisation and outlier removal procedures (77). We removed 21 further individuals as they were the only sample on a slide, preventing their adjustment for slide effects. This left us with 849 DNAm profiles at birth, 910 at 7 years, and 921 at 15-17 years. Data was normalised with functional normalisation (78) across all timepoints.

In BiB, DNAm profiles were assessed using using the Illumina Infinium MethylationEPIC beadchip arrays (Illumina, San Diego, CA, USA). Quality control and normalisation procedures are described in Supplemental Methods. For this study we used DNAm data of 951 children; 88 further participants were removed as they were related >12.5%. DNAm was normalised using the Functional Normalization algorithm (78) implemented in meffil (77).

### Adjusting DNAm data for known covariates

Blood cell count proportion estimates were generated using the Houseman algorithm implemented in the R package meffil v0.1.0 (77). Reference panels were as follows: ARIES at birth (Bcell, CD4T, CD8T, CD14, NK, Gran) (79), ARIES at 7 and 15-17 years (Bcell, CD4T, CD8T, Mono, NK, Gran) (80) and both BiB ethnic groups (Bcell, CD4T, CD8T, Mono, NK, Gran, nRBC) (81).

Outlying methylation values (>10 standard deviations from the probe mean) were removed and replaced with the probe mean.

DNAm data were adjusted for sex, age (apart from the birth timepoints), blood sample type (blood spots, white cells, or whole blood) if more than one was used (as was the case for birth and 7 years in ARIES), Beadchip (also referred to as slide) to represent batch effects, and blood cell count proportion estimates. Sites measured by sub-optimal probes (82) were removed, as were multi-mapping probes and probes targeting non CpG sites that failed liftover to hg19 (56).

### Genotype data generation

ARIES participants were genotyped as part of the main ALSPAC study. All ALSPAC child participants were genotyped with the Illumina HumanHap550 quad genome-wide SNP array (Illumina Inc., San Diego, CA) by the Laboratory Corporation of America (LCA, Burlington, NC, USA) and the Wellcome Trust Sanger Institute (WTSI, Cambridge, UK), supported by 23andMe (76); exclusions and imputation procedures are detailed in the supplementary methods.

BiB participants were genotyped using either the Illumina HumanCoreExome Exome-24 v1.1 microarray, or the Infinium global screen-24+v1.0 array. GenomeStudio 2011.1 was used to pre-process samples; exclusions and imputation procedures are detailed in the supplementary methods.

### Correlation of all sites on the 450k array

DNAm sites were correlated using the biweight mid-correlation (83–86), a median-based method. As Pearson correlation is mean based it may not be suitable for DNAm data, which may be influenced by genotype and so form clusters. In ARIES, a correlation matrix of 394,842 x 394,842 yields 77,949,905,061 unique correlations when we remove the diagonal of the matrix. In BiB there were 68,374,355,910 unique correlations. Due to size limits in R it is not possible to create a single matrix containing all pairwise correlations. As a solution, the DNAm sites were split into blocks of 25,000, and all blocks were correlated against each other. To assess the features of correlating pairs, the correlations were then split by value, from −1 to 1, in increments of 0.1. This enabled analysis of the features of all correlations of different strengths (ie, do high correlations differ from low correlations?). The process is summarised in Figure 11 and code is available here: (https://github.com/shwatkins/PhD/tree/master/450k_correlation_analysis)

**Figure 11:**
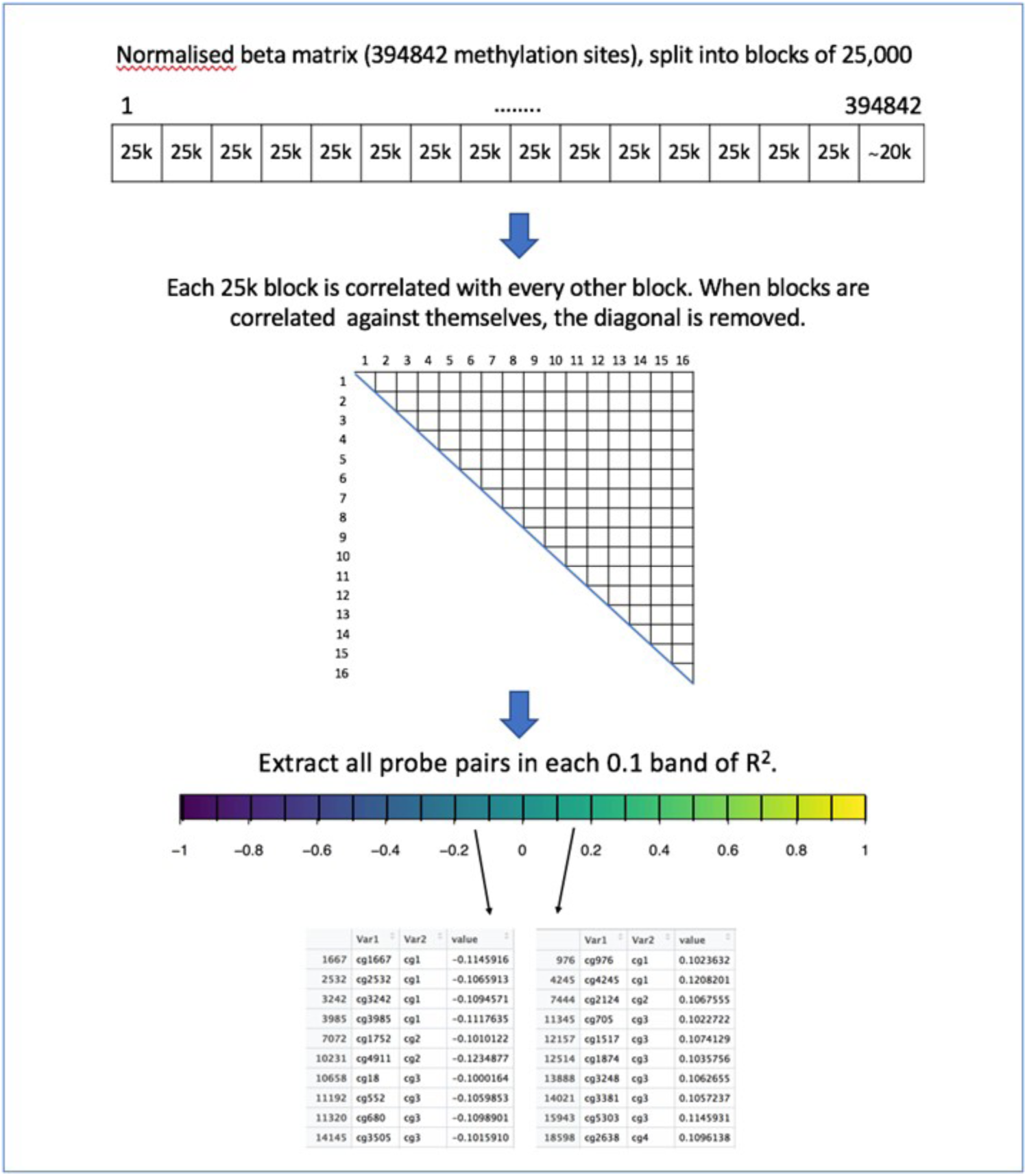
Overview of creating the correlation matrix and extracting pairwise correlations (numbers are representative of the correlation matrix in ARIES).

### Estimation of decay of cis correlations

To assess the decay of the *cis* correlation structure, decay plots of *cis* correlations were created across all chromosomes, and for each chromosome separately based on the method used in (32). All correlations where the DNAm sites on the same chromosome were within 10kb were extracted from the data. As we hypothesised that negative correlations may not have the same structure as positive *cis* correlations, positive and negative correlations were separated. The pairwise correlations were then binned; to display all chromosomes, specifying at least 4000 pairwise correlations per bin for positive correlations into between 469 and 578 bins, and 1000 pairwise correlations for negative correlations into between 800 and 827 bins. Once the correlations were grouped into bins, the mean pairwise correlation for each bin, the standard deviation of the correlation values in each bin, and the median pairwise distance between the correlating pairs of DNAm sites per bin, were calculated (supplementary figure 2).

### Preservation of correlations across datasets

Mean difference plots were used to assess the preservation of high correlations between ARIES and BiB, and between the two ethnic groups in BiB. For each pair of DNAm sites, the mean of the correlation of the two groups is plotted against the difference in correlation between the two groups. 95% confidence intervals are calculated using the difference, illustrating confidence in the mean difference estimates (87).

### Heritability and environmental contributions to co-methylation

To assess the impact of heritability on DNAm correlations, the estimates of heritability and environmental influences on DNAm created by (1) were used to estimate the proportion of sites in each correlation band that were influenced by genetic, unique environmental, and shared environmental factors. The contribution of these influences were assembled for a unique list of all DNAm sites which featured in each correlation range (−1 to 1, in increments of 0.1).

### Overlap of mQTLs with strength of co-methylation

We identified whether, for each correlating pair, neither, one or both of the DNAm sites were associated with an mQTL. For each correlating pair, each DNAm site was assigned 0 if it was not associated with any SNPs in the GoDMC dataset (55), and a 1 if there were one or more SNP associations. The value was summed for the two sites in a correlating pair, which resulted in 0 if neither DNAm site was associated with a SNP, 1 if only one of the DNAm sites was associated with a SNP, and 2 if both DNAm sites were associated with a SNP. This was done separately for *cis* and *trans* correlations, and totalled over all pairs in each correlation range, to illustrate the distribution of mQTLs across values of correlation. Please note that this does not identify whether both DNAm sites are associated with the same mQTL.

### Removing mQTL influence from cis correlations

To illustrate some of the impact mQTLs have on co-methylation, we adjusted the *cis* correlation decay plot for the strongest *cis* mQTL associated with each DNAm site, thereby removing the strongest single genetic influence on DNAm correlations. For this analysis, we used chromosome 10 as an example. The analysis was performed adapting GoDMC analysis scripts (https://github.com/MRCIEU/godmc). This analysis uses an allele count file and a SNP frequency file created through plink2 (88) to adjust the DNAm matrix for the strongest *cis*-mQTL for each DNAm site, using an additive model. The residuals were then taken forward to calculate the decay of the correlations. The DNAm values which did not have an associated mQTL were not adjusted and were not included in the *cis* decay plot.

### Enrichment analyses of strongly co-methylated sites

To identify whether DNAm sites which form strong correlations overlap with genomic sites of interest, we used the locus overlap package LOLA version 1.10.0 (89). LOLA assesses enrichment based on genomic regions rather than genes. A list of unique DNAm sites correlating >0.9 formed the test dataset; all sites in the analysis formed the background. Because binding of transcription factors is enriched in GC-rich areas of the genome, the content of the background set was reduced and matched to the GC content of the test set using frequency quantiles. Genomic locations of the DNAm sites were taken from the IlluminaHumanMethylation450kanno.ilmn12.hg19 R package version 0.6.0 (90). Start and end sites were computed as −500bp and +500bp from the DNAm site position, respectively. A radius of 1kb was thought to be appropriate overlap because DNAm sites within 1-2kb are highly correlated. We used region sets created by the LOLA team (available through http://lolaweb.databio.org) - the ENCODE transcription factor binding sites (http://hgdownload.cse.ucsc.edu/goldenPath/hg19/encodeDCC/wgEncodeAwgTfbsUniform/), chromHMM imputed 25 chromatin states from Roadmap Epigenomics (https://egg2.wustl.edu/roadmap/data/byFileType/chromhmmSegmentations/ChmmModels/imputed12marks/jointModel/final/) (17, 91), and gene annotations from https://zwdzwd.github.io/InfiniumAnnotation.

### Assessing *trans* correlations for chromatin contacts

Previous work (16) has illustrated that inter-chromosomal regions which connect have correlated DNAm states. To test whether highly correlating regions are enriched for chromatin contacts, we adapted the GoDMC analysis pipeline (https://github.com/MRCIEU/godmc_phase2_analysis/tree/master/13_hi-c) to test for chromatin contact enrichment, using a publicly available chromatin contacts map (92). For each of the pairwise contacts in the dataset (92) a 1kb region for the two contacting areas of the genome was generated. These are split into files containing all interactions for all possible pairs of chromosomes (for example all contacts between chromosome 1 and chromosome 18). This resulted in 231 files containing all inter-chromosomal contacts on the autosomes. Next, the inter-chromosomal correlations r>0.9 in the relevant dataset (e.g. ARIES 7 year olds) had a 500bp region defined either side of the DNAm sites. We identified which correlating pairs overlapped with the contact regions (92).

To ascertain whether there were more contacts in our data than expected by chance we created a permuted dataset, where from the original data the second DNAm site in the highly correlating pair is replaced randomly with another in the dataset. Broken pairs that match a pair from the original dataset were removed, as are duplicates to avoid double counting. Then overlaps with the HiC data (92) were calculated for the permuted data. This process was repeated 1,000 times, the permuted datasets were merged together, and a distribution of overlap counts was created for the permuted data. This distribution was used to create a p value for the overlap of the permuted distribution chromatin contact overlaps with the number of overlaps in the real data.

## Declarations

### Ethics approval and consent to participate

Ethical approval for the ALSPAC study was obtained from the ALSPAC Ethics and Law Committee and the Local Research Ethics Committees, http://www.bristol.ac.uk/alspac/researchers/research-ethics/, under proposal number B2808. Informed consent for the use of data collected via questionnaires and clinics was obtained from participants following the recommendations of the ALSPAC Ethics and Law Committee at the time. Ethical approval for the BiB portion of this study was granted by the Bradford Research Ethics Committee (Ref 07/H1302/112). Written informed consent was obtained from the mothers (for themselves and their children) when they registered for the study. Consent for all biological samples has been collected in accordance with the Human Tissue Act (2004).

### Consent for publication

Not applicable

### Availability of data and materials

Data are available to researchers by request from the Avon Longitudinal Study of Parents and Children Executive Committee (http://www.bristol.ac.uk/alspac/researchers/data-access/) as outlined in the study’s access policy http://www.bristol.ac.uk/media-library/sites/alspac/documents/researchers/data-access/ALSPAC_Access_Policy.pdf. ALSPAC fully supports Wellcome and the RCUK policies on open access. The ALSPAC study website contains details of all the data that are available through a fully searchable data dictionary and variable search tool: http://www.bristol.ac.uk/alspac/researchers/our-data/.

BiB data are available to researchers who submit an expression of interest to the Born in Bradford Executive Group who review applications monthly and aim to respond within eight weeks. Data requests will require a formal Data Transfer Agreement, and data are not publicly available due to the terms of the ethical approval. More details of data available and how to apply for access on the Born in Bradford website: https://borninbradford.nhs.uk/research/.

Scripts to replicate these analyses can be found on GitHub: https://github.com/shwatkins/PhD/tree/master/450k_correlation_analysis

The full DNAm correlation matrices can be found at https://data-bris.acrc.bris.ac.uk/deposits/31uze72mt042g2ticr0w6z6v8y, DOI = 10.5523/bris.31uze72mt042g2ticr0w6z6v8y

## Acknowledgements

We are extremely grateful to all the families who took part in the ALSPAC study, the midwives for their help in recruiting them, and the whole ALSPAC team, which includes interviewers, computer and laboratory technicians, clerical workers, research scientists, volunteers, managers, receptionists and nurses. 450k DNAm array data and part of the genotype data was generated in the Bristol Bioresource Laboratory Illumina Facility, University of Bristol.

Born in Bradford is only possible because of the enthusiasm and commitment of the Children and Parents in BiB. We are grateful to all the participants, practitioners and researchers who have made Born in Bradford happen. 450k DNAm and genotype array data was generated in the Bristol Bioresource Laboratory Illumina Facility, University of Bristol.

## Author contributions

Designed individual studies and contributed data: TRG, JLM, DAL

Designed and managed the study: JLM, NJT, TRG

Designed the analyses: SHW, JLM, TRG, NJT, MS, GH, KB

Conducted analyses: SHW

Critically reviewed and revised the analyses: SHW, JLM, TRG, NJT, MS, GH, KB, DAL

SHW wrote the manuscript; all authors reviewed and revised the manuscript

## References

1. Hannon E, Knox O, Sugden K, Burrage J, Wong CCY, Belsky DW, et al. Characterizing genetic and environmental influences on variable DNA methylation using monozygotic and dizygotic twins. PLoS Genet. 2018;14(8):e1007544.

2. Castillo-Fernandez JE, Spector TD, Bell JT. Epigenetics of discordant monozygotic twins: implications for disease. Genome Med. 2014;6(7):60.

3. Bell JT, Pai AA, Pickrell JK, Gaffney DJ, Pique-Regi R, Degner JF, et al. DNA methylation patterns associate with genetic and gene expression variation in HapMap cell lines (vol 12, pg R10, 2011). Genome Biol. 2011;12(6).

4. Gutierrez-Arcelus M, Lappalainen T, Montgomery SB, Buil A, Ongen H, Yurovsky A, et al. Passive and active DNA methylation and the interplay with genetic variation in gene regulation. Elife. 2013;2:e00523.

5. Saunderson EA, Stepper P, Gomm JJ, Hoa L, Morgan A, Allen MD, et al. Hit-and-run epigenetic editing prevents senescence entry in primary breast cells from healthy donors. Nat Commun. 2017;8(1):1450.

6. Maeder ML, Angstman JF, Richardson ME, Linder SJ, Cascio VM, Tsai SQ, et al. Targeted DNA demethylation and activation of endogenous genes using programmable TALE-TET1 fusion proteins. Nat Biotechnol. 2013;31(12):1137–42.

7. Bonder MJ, Luijk R, Zhernakova DV, Moed M, Deelen P, Vermaat M, et al. Disease variants alter transcription factor levels and methylation of their binding sites. Nat Genet. 2017;49(1):131–8.

8. Razin A, Riggs AD. DNA methylation and gene function. Science. 1980;210(4470):604–10.

9. Prendergast GC, Ziff EB. Methylation-sensitive sequence-specific DNA binding by the c-Myc basic region. Science. 1991;251(4990):186–9.

10. Kribelbauer JF, Lu XJ, Rohs R, Mann RS, Bussemaker HJ. Toward a Mechanistic Understanding of DNA Methylation Readout by Transcription Factors. J Mol Biol. 2019.

11. Domcke S, Bardet AF, Adrian Ginno P, Hartl D, Burger L, Schubeler D. Competition between DNA methylation and transcription factors determines binding of NRF1. Nature. 2015;528(7583):575–9.

12. Yin Y, Morgunova E, Jolma A, Kaasinen E, Sahu B, Khund-Sayeed S, et al. Impact of cytosine methylation on DNA binding specificities of human transcription factors. Science. 2017;356(6337).

13. Song G, Wang G, Luo X, Cheng Y, Song Q, Wan J, et al. An all-to-all approach to the identification of sequence-specific readers for epigenetic DNA modifications on cytosine. Nat Commun. 2021;12(1):795.

14. Ciccone DN, Su H, Hevi S, Gay F, Lei H, Bajko J, et al. KDM1B is a histone H3K4 demethylase required to establish maternal genomic imprints. Nature. 2009;461(7262):415–8.

15. Smith ZD, Meissner A. DNA methylation: roles in mammalian development. Nat Rev Genet. 2013;14(3):204–20.

16. Li G, Liu Y, Zhang Y, Kubo N, Yu M, Fang R, et al. Joint profiling of DNA methylation and chromatin architecture in single cells. Nat Methods. 2019;16(10):991–3.

17. Roadmap Epigenomics C, Kundaje A, Meuleman W, Ernst J, Bilenky M, Yen A, et al. Integrative analysis of 111 reference human epigenomes. Nature. 2015;518(7539):317–30.

18. Richmond RC, Simpkin AJ, Woodward G, Gaunt TR, Lyttleton O, McArdle WL, et al. Prenatal exposure to maternal smoking and offspring DNA methylation across the lifecourse: findings from the Avon Longitudinal Study of Parents and Children (ALSPAC). Hum Mol Genet. 2015;24(8):2201–17.

19. Smith JA, Zhao W, Wang X, Ratliff SM, Mukherjee B, Kardia SLR, et al. Neighborhood characteristics influence DNA methylation of genes involved in stress response and inflammation: The Multi-Ethnic Study of Atherosclerosis. Epigenetics. 2017;12(8):662–73.

20. de Vocht F, Suderman M, Ruano-Ravina A, Thomas R, Wakeford R, Relton C, et al. Residential exposure to radon and DNA methylation across the lifecourse: an exploratory study in the ALSPAC birth cohort. Wellcome Open Res. 2019;4:3.

21. Liu J, Carnero-Montoro E, van Dongen J, Lent S, Nedeljkovic I, Ligthart S, et al. An integrative cross-omics analysis of DNA methylation sites of glucose and insulin homeostasis. Nature Communications. 2019;10.

22. Xu CJ, Soderhall C, Bustamante M, Baiz N, Gruzieva O, Gehring U, et al. DNA methylation in childhood asthma: an epigenome-wide meta-analysis. Lancet Respir Med. 2018;6(5):379–88.

23. Battram T, Yousefi P, Crawford G, Prince C, Babei MS, Sharp G, et al. The EWAS Catalog: a database of epigenome-wide association studies. OSF Preprints. 2021.

24. Langfelder P, Cantle JP, Chatzopoulou D, Wang N, Gao F, Al-Ramahi I, et al. Integrated genomics and proteomics define huntingtin CAG length-dependent networks in mice. Nat Neurosci. 2016;19(4):623–33.

25. Pidsley R, Viana J, Hannon E, Spiers H, Troakes C, Al-Saraj S, et al. Methylomic profiling of human brain tissue supports a neurodevelopmental origin for schizophrenia. Genome Biol. 2014;15(10):483.

26. Wong CCY, Smith RG, Hannon E, Ramaswami G, Parikshak NN, Assary E, et al. Genome-wide DNA methylation profiling identifies convergent molecular signatures associated with idiopathic and syndromic autism in post-mortem human brain tissue. Hum Mol Genet. 2019;28(13):2201–11.

27. Swarup V, Hinz FI, Rexach JE, Noguchi K, Toyoshiba H, Oda A, et al. Identification of evolutionarily conserved gene networks mediating neurodegenerative dementia. Nat Med. 2019;25(1):152-+.

28. Eckhardt F, Lewin J, Cortese R, Rakyan VK, Attwood J, Burger M, et al. DNA methylation profiling of human chromosomes 6, 20 and 22. Nat Genet. 2006;38(12):1378–85.

29. Kuan PF, Chiang DY. Integrating prior knowledge in multiple testing under dependence with applications to detecting differential DNA methylation. Biometrics. 2012;68(3):774–83.

30. Liu Y, Li X, Aryee MJ, Ekstrom TJ, Padyukov L, Klareskog L, et al. GeMes, clusters of DNA methylation under genetic control, can inform genetic and epigenetic analysis of disease. Am J Hum Genet. 2014;94(4):485–95.

31. Ong ML, Holbrook JD. Novel region discovery method for Infinium 450K DNA methylation data reveals changes associated with aging in muscle and neuronal pathways. Aging Cell. 2014;13(1):142–55.

32. Saffari A, Silver MJ, Zavattari P, Moi L, Columbano A, Meaburn EL, et al. Estimation of a significance threshold for epigenome-wide association studies. Genet Epidemiol. 2018;42(1):20–33.

33. Shoemaker R, Deng J, Wang W, Zhang K. Allele-specific methylation is prevalent and is contributed by CpG-SNPs in the human genome. Genome Res. 2010;20(7):883–9.

34. Zhang W, Spector TD, Deloukas P, Bell JT, Engelhardt BE. Predicting genome-wide DNA methylation using methylation marks, genomic position, and DNA regulatory elements. Genome Biol. 2015;16:14.

35. Garg P, Joshi RS, Watson C, Sharp AJ. A survey of inter-individual variation in DNA methylation identifies environmentally responsive co-regulated networks of epigenetic variation in the human genome. PLoS Genet. 2018;14(10):e1007707.

36. Wu Y, Qi T, Wang H, Zhang F, Zheng Z, Phillips-Cremins JE, et al. Promoter-anchored chromatin interactions predicted from genetic analysis of epigenomic data. Nat Commun. 2020;11(1):2061.

37. Shah S, McRae AF, Marioni RE, Harris SE, Gibson J, Henders AK, et al. Genetic and environmental exposures constrain epigenetic drift over the human life course. Genome Res. 2014;24(11):1725–33.

38. van Dongen J, Nivard MG, Willemsen G, Hottenga JJ, Helmer Q, Dolan CV, et al. Genetic and environmental influences interact with age and sex in shaping the human methylome. Nat Commun. 2016;7:11115.

39. Sugden K, Hannon EJ, Arseneault L, Belsky DW, Corcoran DL, Fisher HL, et al. Patterns of Reliability: Assessing the Reproducibility and Integrity of DNA Methylation Measurement. Patterns (N Y). 2020;1(2):100014.

40. Hannum G, Guinney J, Zhao L, Zhang L, Hughes G, Sadda S, et al. Genome-wide methylation profiles reveal quantitative views of human aging rates. Mol Cell. 2013;49(2):359–67.

41. Horvath S. DNA methylation age of human tissues and cell types. Genome Biol. 2013;14(10).

42. Teschendorff AE, West J, Beck S. Age-associated epigenetic drift: implications, and a case of epigenetic thrift? Hum Mol Genet. 2013;22(R1):R7–R15.

43. Ambatipudi S, Cuenin C, Hernandez-Vargas H, Ghantous A, Le Calvez-Kelm F, Kaaks R, et al. Tobacco smoking-associated genome-wide DNA methylation changes in the EPIC study. Epigenomics. 2016;8(5):599–618.

44. Houtepen LC, Hardy R, Maddock J, Kuh D, Anderson EL, Relton CL, et al. Childhood adversity and DNA methylation in two population-based cohorts. Transl Psychiatry. 2018;8(1):266.

45. Dunn EC, Soare TW, Zhu Y, Simpkin AJ, Suderman MJ, Klengel T, et al. Sensitive Periods for the Effect of Childhood Adversity on DNA Methylation: Results From a Prospective, Longitudinal Study. Biol Psychiatry. 2019;85(10):838–49.

46. Barcelona de Mendoza V, Huang Y, Crusto CA, Sun YV, Taylor JY. Perceived Racial Discrimination and DNA Methylation Among African American Women in the InterGEN Study. Biol Res Nurs. 2018;20(2):145–52.

47. van der Laan LC, Meeks KAC, Chilunga FP, Agyemang C, Venema A, Mannens M, et al. Epigenome-wide association study for perceived discrimination among sub-Saharan African migrants in Europe - the RODAM study. Sci Rep. 2020;10(1):4919.

48. Needham BL, Smith JA, Zhao W, Wang X, Mukherjee B, Kardia SL, et al. Life course socioeconomic status and DNA methylation in genes related to stress reactivity and inflammation: The multi-ethnic study of atherosclerosis. Epigenetics. 2015;10(10):958–69.

49. Dai L, Mehta A, Mordukhovich I, Just AC, Shen J, Hou L, et al. Differential DNA methylation and PM2.5 species in a 450K epigenome-wide association study. Epigenetics. 2017;12(2):139–48.

50. Baccarelli A, Wright RO, Bollati V, Tarantini L, Litonjua AA, Suh HH, et al. Rapid DNA methylation changes after exposure to traffic particles. Am J Respir Crit Care Med. 2009;179(7):572–8.

51. Tang H. Confronting ethnicity-specific disease risk. Nat Genet. 2006;38(1):13–5.

52. Bibikova M, Barnes B, Tsan C, Ho V, Klotzle B, Le JM, et al. High density DNA methylation array with single CpG site resolution. Genomics. 2011;98(4):288–95.

53. Sandoval J, Heyn H, Moran S, Serra-Musach J, Pujana MA, Bibikova M, et al. Validation of a DNA methylation microarray for 450,000 CpG sites in the human genome. Epigenetics. 2011;6(6):692–702.

54. Genomes Project C, Auton A, Brooks LD, Durbin RM, Garrison EP, Kang HM, et al. A global reference for human genetic variation. Nature. 2015;526(7571):68–74.

55. Min JL, Hemani G, Hannon E, Dekkers KF, Castillo-Fernandez J, Luijk R, et al. Genomic and phenotypic insights from an atlas of genetic effects on DNA methylation. Nat Genet. 2021;53(9):1311–21.

56. Min JL, Hemani G, Hannon E, Dekkers KF, Castillo-Fernandez J, Luijk R, et al. Genomic and phenomic insights from an atlas of genetic effects on DNA methylation. medRxiv. 2020:2020.09.01.20180406.

57. Gaunt TR, Shihab HA, Hemani G, Min JL, Woodward G, Lyttleton O, et al. Systematic identification of genetic influences on methylation across the human life course. Genome Biol. 2016;17:61.

58. Sodini SM, Kemper KE, Wray NR, Trzaskowski M. Comparison of Genotypic and Phenotypic Correlations: Cheverud’s Conjecture in Humans. Genetics. 2018;209(3):941–8.

59. Lienert F, Wirbelauer C, Som I, Dean A, Mohn F, Schubeler D. Identification of genetic elements that autonomously determine DNA methylation states. Nat Genet. 2011;43(11):1091–7.

60. Barabasi AL, Albert R. Emergence of scaling in random networks. Science. 1999;286(5439):509–12.

61. Barabasi AL. Scale-free networks: a decade and beyond. Science. 2009;325(5939):412–3.

62. Broido AD, Clauset A. Scale-free networks are rare. Nat Commun. 2019;10(1):1017.

63. Fraser A, Macdonald-Wallis C, Tilling K, Boyd A, Golding J, Davey Smith G, et al. Cohort Profile: the Avon Longitudinal Study of Parents and Children: ALSPAC mothers cohort. Int J Epidemiol. 2013;42(1):97–110.

64. Boyd A, Golding J, Macleod J, Lawlor DA, Fraser A, Henderson J, et al. Cohort Profile: the ‘children of the 90s’--the index offspring of the Avon Longitudinal Study of Parents and Children. Int J Epidemiol. 2013;42(1):111–27.

65. Turowski TW, Tollervey D. Transcription by RNA polymerase III: insights into mechanism and regulation. Biochem Soc Trans. 2016;44(5):1367–75.

66. Vannini A, Ringel R, Kusser AG, Berninghausen O, Kassavetis GA, Cramer P. Molecular basis of RNA polymerase III transcription repression by Maf1. Cell. 2010;143(1):59–70.

67. Abascal-Palacios G, Ramsay EP, Beuron F, Morris E, Vannini A. Structural basis of RNA polymerase III transcription initiation. Nature. 2018;553(7688):301-+.

68. Maass PG, Barutcu AR, Weiner CL, Rinn JL. Inter-chromosomal Contact Properties in Live-Cell Imaging and in Hi-C. Mol Cell. 2018;69(6):1039–45 e3.

69. Neph S, Stergachis AB, Reynolds A, Sandstrom R, Borenstein E, Stamatoyannopoulos JA. Circuitry and dynamics of human transcription factor regulatory networks. Cell. 2012;150(6):1274–86.

70. Kim S, Dunham MJ, Shendure J. A combination of transcription factors mediates inducible interchromosomal contacts. Elife. 2019;8.

71. Hannon E, Lunnon K, Schalkwyk L, Mill J. Interindividual methylomic variation across blood, cortex, and cerebellum: implications for epigenetic studies of neurological and neuropsychiatric phenotypes. Epigenetics. 2015;10(11):1024–32.

72. Javierre BM, Burren OS, Wilder SP, Kreuzhuber R, Hill SM, Sewitz S, et al. Lineage-Specific Genome Architecture Links Enhancers and Non-coding Disease Variants to Target Gene Promoters. Cell. 2016;167(5):1369–84 e19.

73. Wright J, Small N, Raynor P, Tuffnell D, Bhopal R, Cameron N, et al. Cohort Profile: the Born in Bradford multi-ethnic family cohort study. Int J Epidemiol. 2013;42(4):978–91.

74. Department for Communities and Local Government. The English Indices of Deprivation 2015: Statistical Release 2015 [Available from: https://assets.publishing.service.gov.uk/government/uploads/system/uploads/attachment_data/file/465791/English_Indices_of_Deprivation_2015_-_Statistical_Release.pdf.

75. Smith T, Noble M, Noble S, Wright G, McLennan D, Plunkett E. The English indices of deprivation 2015. London: Department for Communities and Local Government. 2015.

76. Relton CL, Gaunt T, McArdle W, Ho K, Duggirala A, Shihab H, et al. Data Resource Profile: Accessible Resource for Integrated Epigenomic Studies (ARIES). Int J Epidemiol. 2015;44(4):1181–90.

77. Min JL, Hemani G, Davey Smith G, Relton C, Suderman M. Meffil: efficient normalization and analysis of very large DNA methylation datasets. Bioinformatics. 2018;34(23):3983–9.

78. Fortin JP, Labbe A, Lemire M, Zanke BW, Hudson TJ, Fertig EJ, et al. Functional normalization of 450k methylation array data improves replication in large cancer studies. Genome Biol. 2014;15(12):503.

79. Gervin K, Page CM, Aass HC, Jansen MA, Fjeldstad HE, Andreassen BK, et al. Cell type specific DNA methylation in cord blood: A 450K-reference data set and cell count-based validation of estimated cell type composition. Epigenetics. 2016;11(9):690–8.

80. Reinius LE, Acevedo N, Joerink M, Pershagen G, Dahlen SE, Greco D, et al. Differential DNA methylation in purified human blood cells: implications for cell lineage and studies on disease susceptibility. PLoS One. 2012;7(7):e41361.

81. Bakulski KM, Feinberg JI, Andrews SV, Yang J, Brown S, S LM, et al. DNA methylation of cord blood cell types: Applications for mixed cell birth studies. Epigenetics. 2016;11(5):354–62.

82. Zhou W, Laird PW, Shen H. Comprehensive characterization, annotation and innovative use of Infinium DNA methylation BeadChip probes. Nucleic Acids Res. 2017;45(4):e22.

83. Langfelder P, Horvath S. WGCNA: an R package for weighted correlation network analysis. BMC Bioinformatics. 2008;9:559.

84. Song L, Langfelder P, Horvath S. Comparison of co-expression measures: mutual information, correlation, and model based indices. BMC Bioinformatics. 2012;13:328.

85. Langfelder P, Horvath S. Fast R Functions for Robust Correlations and Hierarchical Clustering. J Stat Softw. 2012;46(11).

86. Hardin J, Mitani A, Hicks L, VanKoten B. A robust measure of correlation between two genes on a microarray. BMC Bioinformatics. 2007;8:220.

87. Ritchie ME, Phipson B, Wu D, Hu Y, Law CW, Shi W, et al. limma powers differential expression analyses for RNA-sequencing and microarray studies. Nucleic Acids Res. 2015;43(7):e47.

88. Chang CC, Chow CC, Tellier LC, Vattikuti S, Purcell SM, Lee JJ. Second-generation PLINK: rising to the challenge of larger and richer datasets. Gigascience. 2015;4:7.

89. Sheffield NC, Bock C. LOLA: enrichment analysis for genomic region sets and regulatory elements in R and Bioconductor. Bioinformatics. 2016;32(4):587–9.

90. Hansen KD. IlluminaHumanMethylation450kanno.ilmn12.hg19: Annotation for Illumina’s 450k methylation arrays. 0.6.0 ed2016. p. R package.

91. Ernst J, Kellis M. Large-scale imputation of epigenomic datasets for systematic annotation of diverse human tissues. Nat Biotechnol. 2015;33(4):364–76.

92. Rao SS, Huntley MH, Durand NC, Stamenova EK, Bochkov ID, Robinson JT, et al. A 3D map of the human genome at kilobase resolution reveals principles of chromatin looping. Cell. 2014;159(7):1665–80.

