## Supplementary figures for "DNA co-methylation has a stable structure and is related to specific aspects of genome regulation"

*Figure 1: **Correlation distribution plots**: Separate distribution of cis and trans correlation values for all five datasets* ........ 2

*Figure 2: **Cis decay plots**: Decay plots of cis correlations from all filtered sites on the 450k array across all autosomes for each of the five datasets. The variance in each bin was added to the plot to demonstrate the uncertainty around the binned estimates. Decay is virtually identical across all datasets.* ........ 3

*Figure 3: **Mean difference plots**: Mean difference plots, which plot the mean against the difference in correlation, for correlations >0.8 in each dataset. Cis (left plots) and trans (right plots) between: ARIES at birth and 7 years (top); ARIES at 7 and 15-17 years (2nd row); ARIES and BiB white British individuals (third row); and the BiB white British and Pakistani racial/ethnic groups (4th row). Solid black line represents a difference in correlation of 0; green dashed lines represent 95% confidence intervals.* ........ 4

*Figure 8: **Circos plots** visualising trans correlations r>0.9 in ARIES at birth (top left), 7 years (top right) and 15-17 years (middle), and in BiB in the white British group (bottom left) and the Pakistani group (bottom right)..* 10

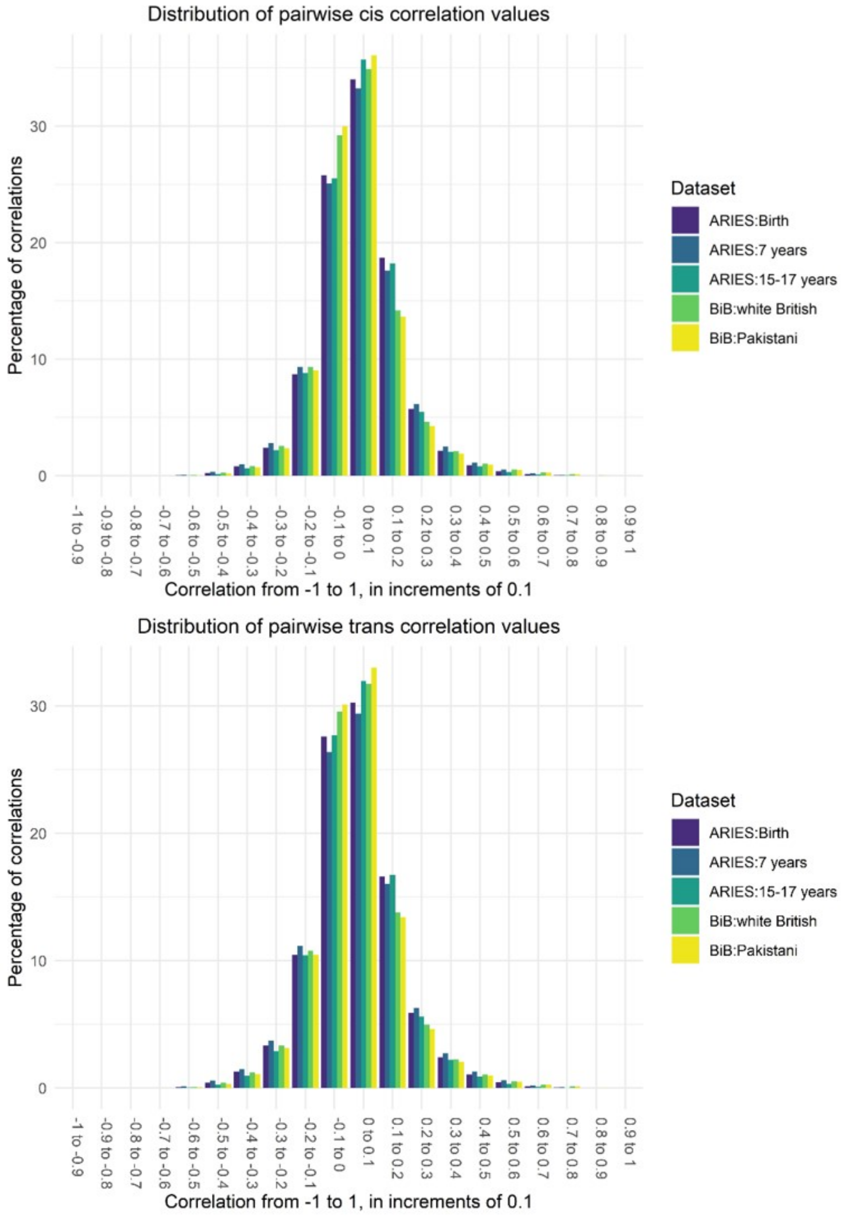

*Figure 1:* ***Correlation distribution plots****: Separate distribution of cis and trans correlation values for all five datasets*

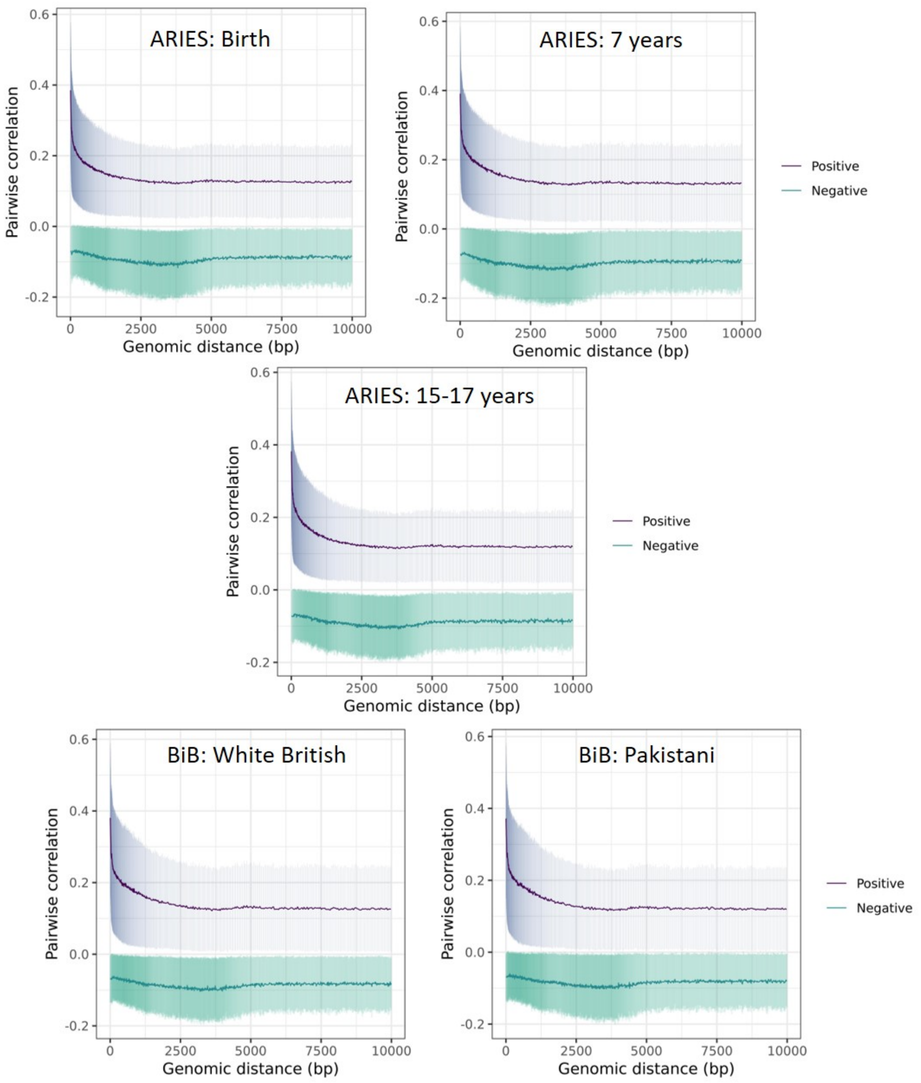

*Figure 2:* ***Cis decay plots****: Decay plots of cis correlations from all filtered sites on the 450k array across all autosomes for each of the five datasets. The variance in each bin was added to the plot to demonstrate the uncertainty around the binned estimates. Decay is virtually identical across all datasets.*

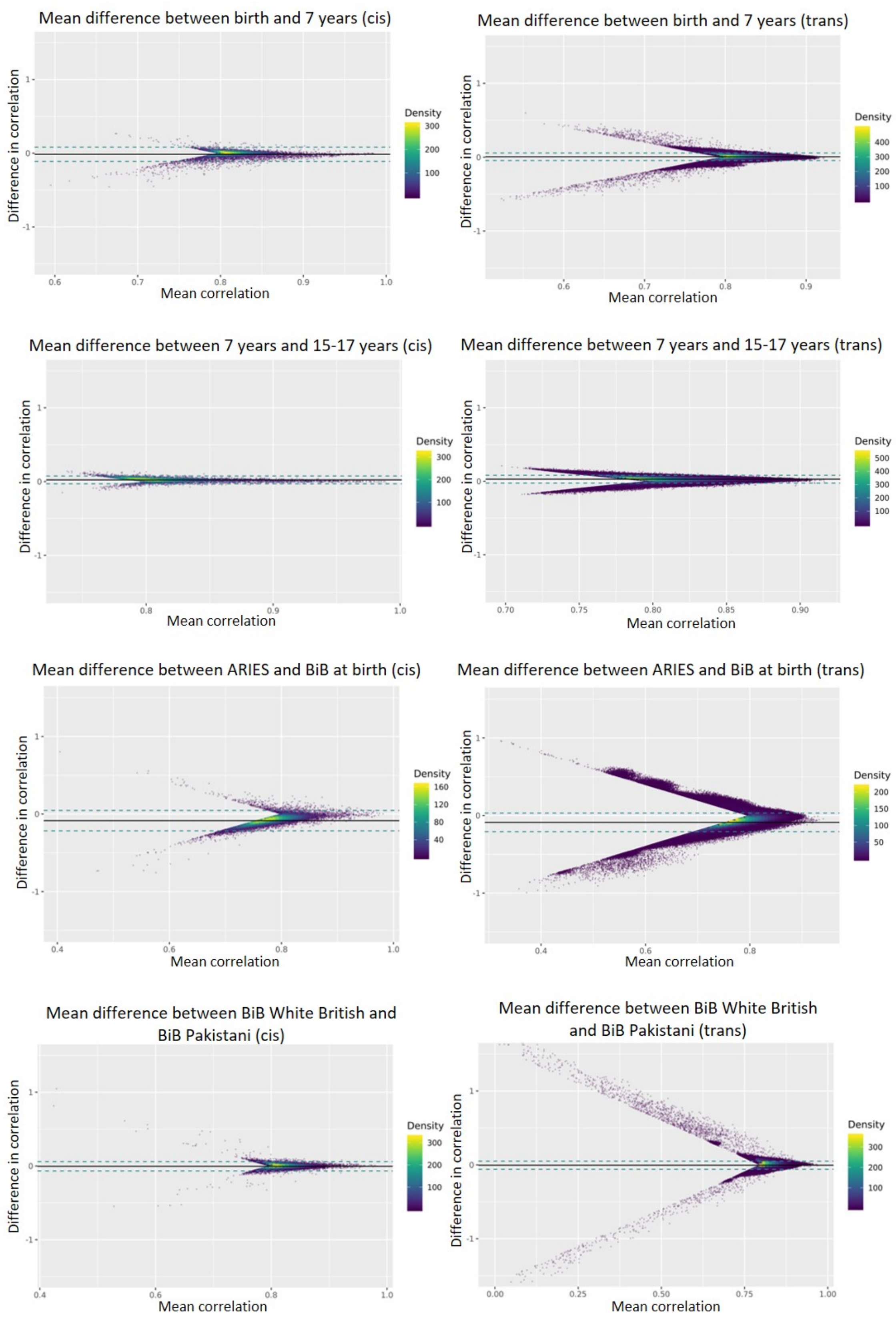

*Figure 3: **Mean difference plots**: Mean difference plots, which plot the mean against the difference in correlation, for correlations >0.8 in each dataset. Cis (left plots) and trans (right plots) between: ARIES at birth and 7 years (top); ARIES at 7 and 15-17 years (2nd row); ARIES and BiB white British individuals (third row); and the BiB white British and Pakistani racial/ethnic groups (4th row). Solid black line represents a difference in correlation of 0; green dashed lines represent 95% confidence intervals.*

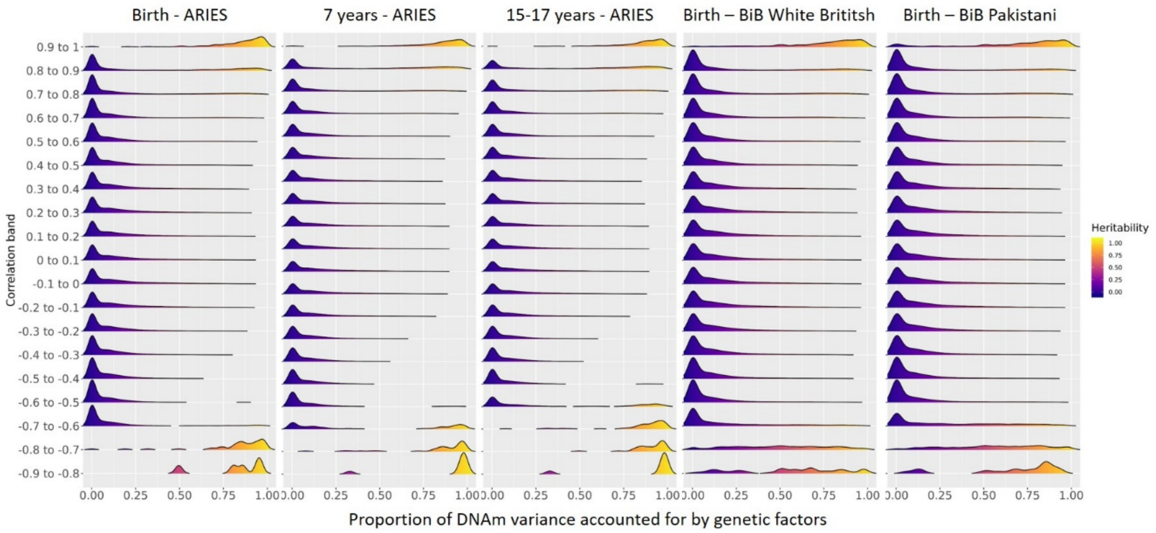

*Figure 4A:* ***Heritability: Cis correlations****. Density plots illustrating the proportion of DNAm variation due to heritability for DNAm sites with correlations of differing strengths, for cis correlations, in each of the five datasets*

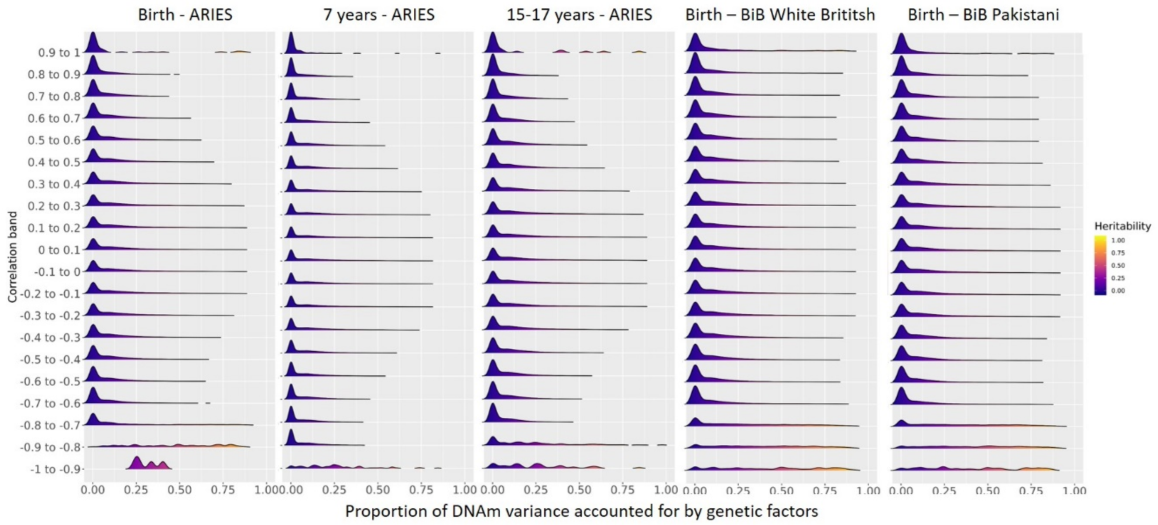

*Figure 4B:* ***Heritability: Trans correlations****. Density plots illustrating the proportion of DNAm variation due to heritability for DNAm sites with correlations of differing strengths, for trans correlations, in each of the five datasets*

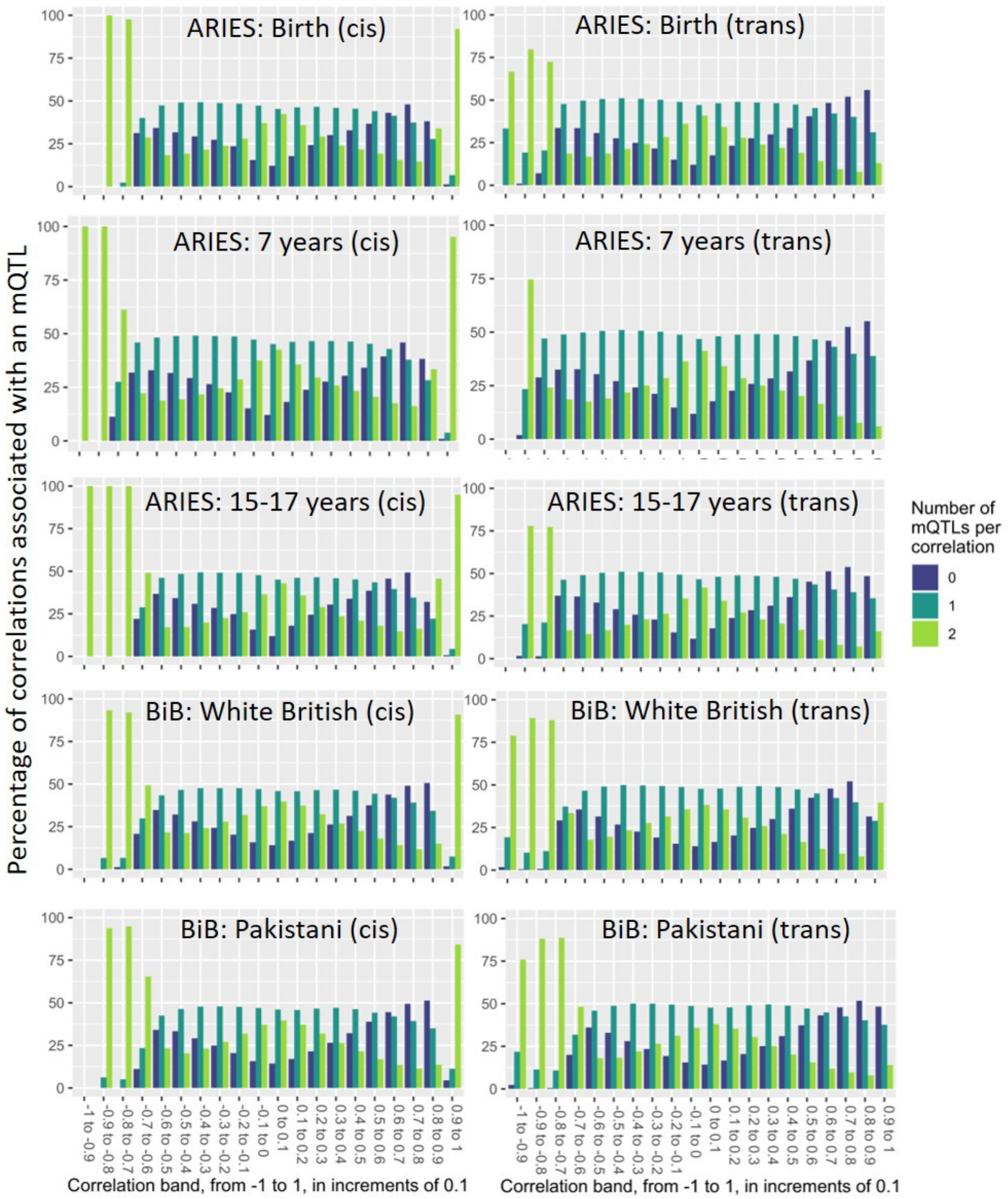

*Figure 5: **mQTL plots**: Bar plots of the percentage of pairwise correlations in each correlation range that have 0, 1 or 2 DNAm sites associated with an mQTL identified by the GoDMC consortium. Split by cis (left) and trans (right) correlating pairs, In ARIES at birth (1st row), 7 years (2nd row) and 15-17 years (3rd row), and in BiB white British (4th row) and Pakistani (5th row) individuals.*

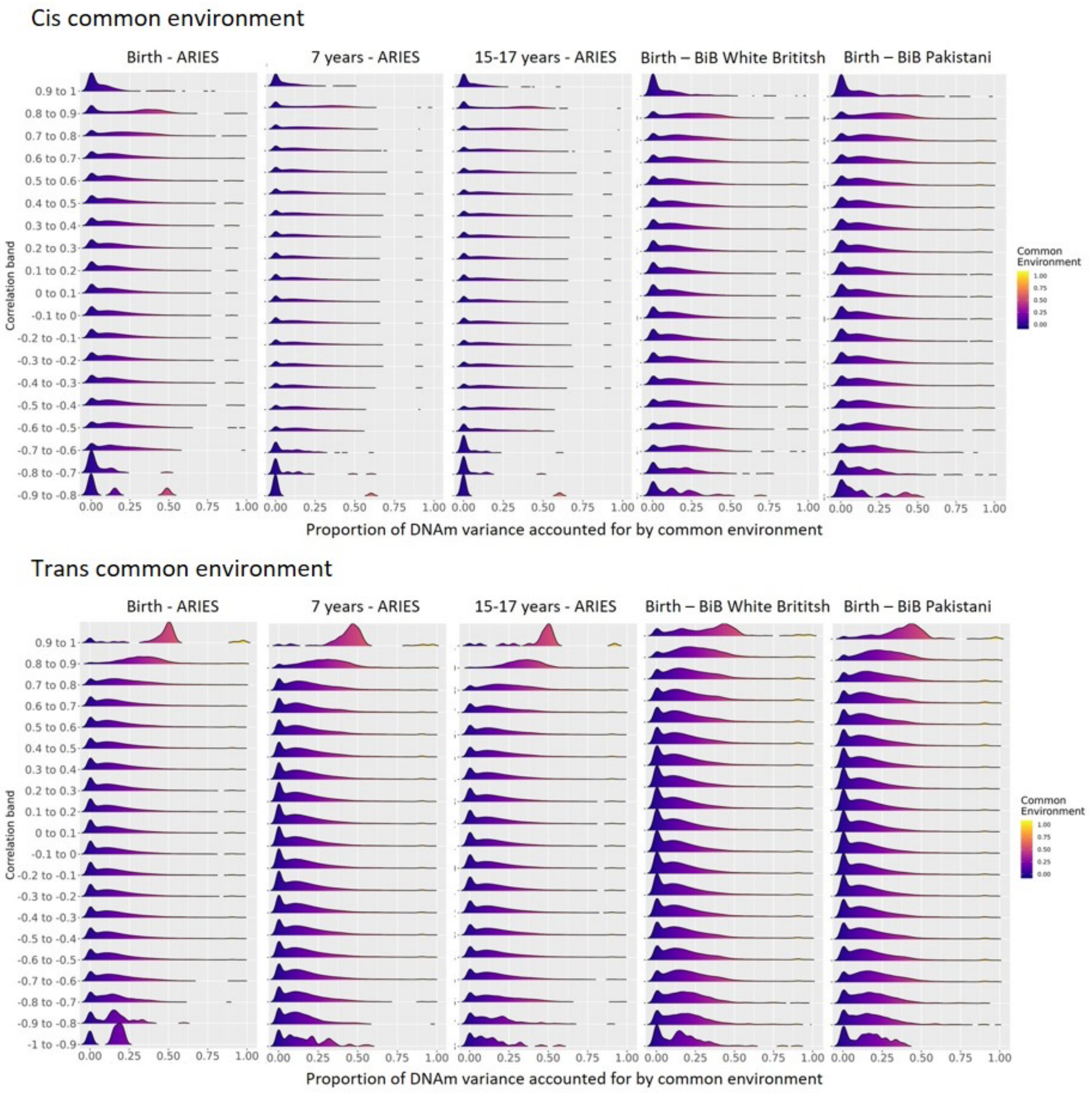

*Figure 6A:* ***Common environment plots****: Density plots illustrating the proportion of DNAm variation due to common environment for DNAm sites with correlations of differing strengths, in each of the five datasets*

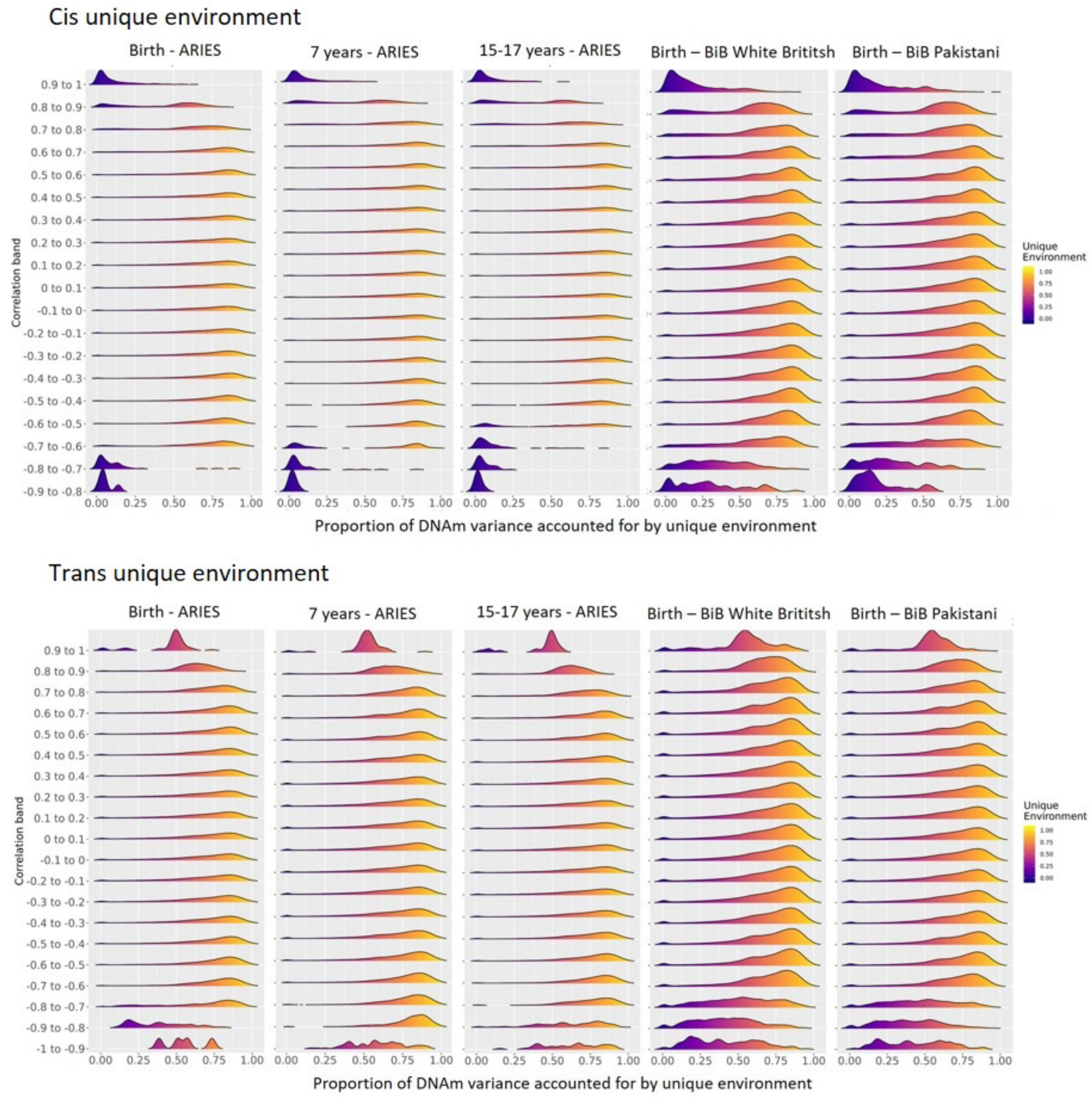

*Figure 6B: **Unique environment plots**: Density plots illustrating the proportion of DNAm variation due to unique environment for DNAm sites with correlations of differing strengths, in each of the five datasets*

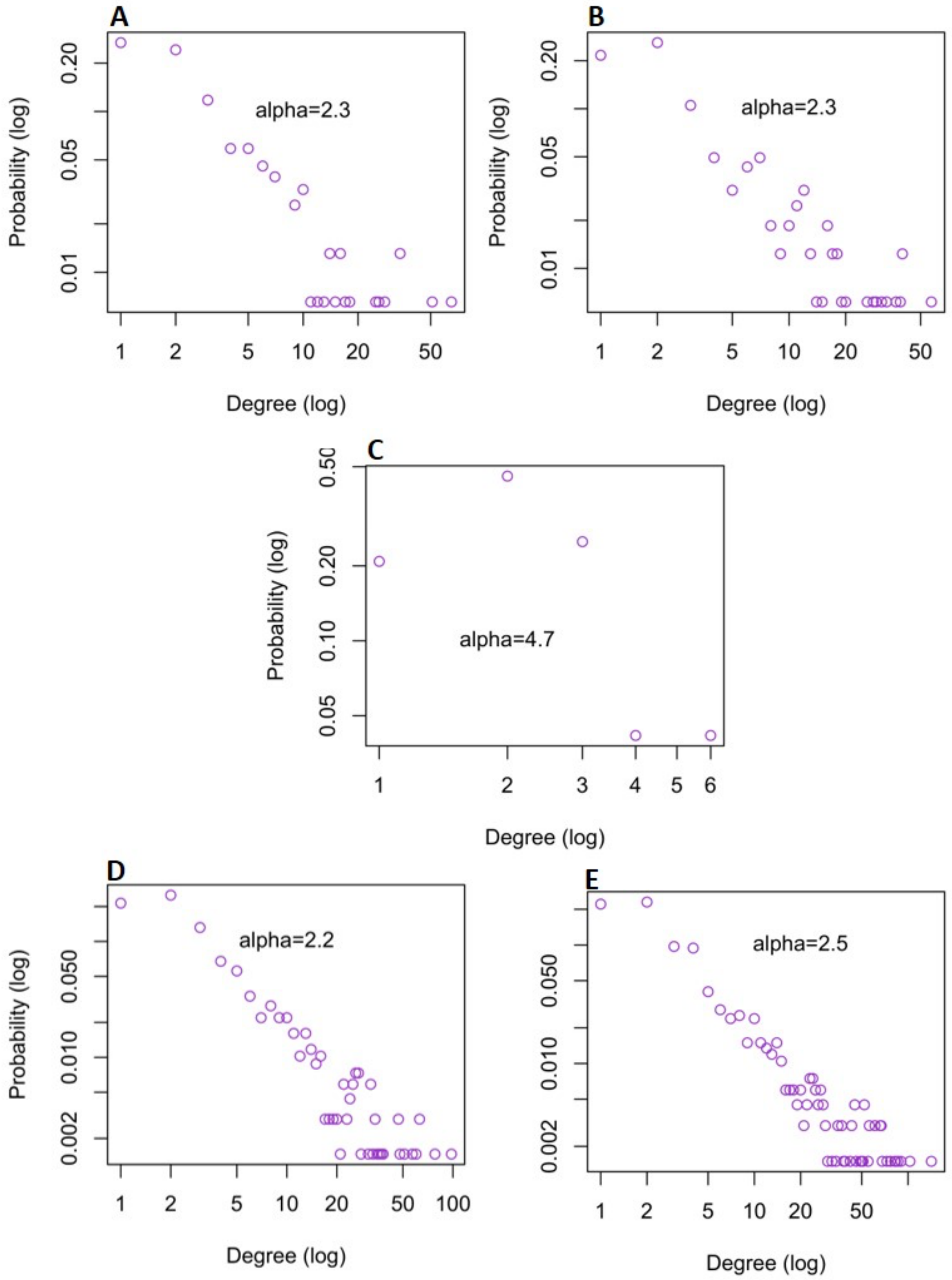

*Figure 7:* ***Degree distribution plots*** *for* ***A*** *Aries at birth* ***B*** *ARIES at 7 years* ***C*** *ARIES at 15-17 years* ***D*** *BiB white British participants and* ***E*** *BiB Pakistani participants*

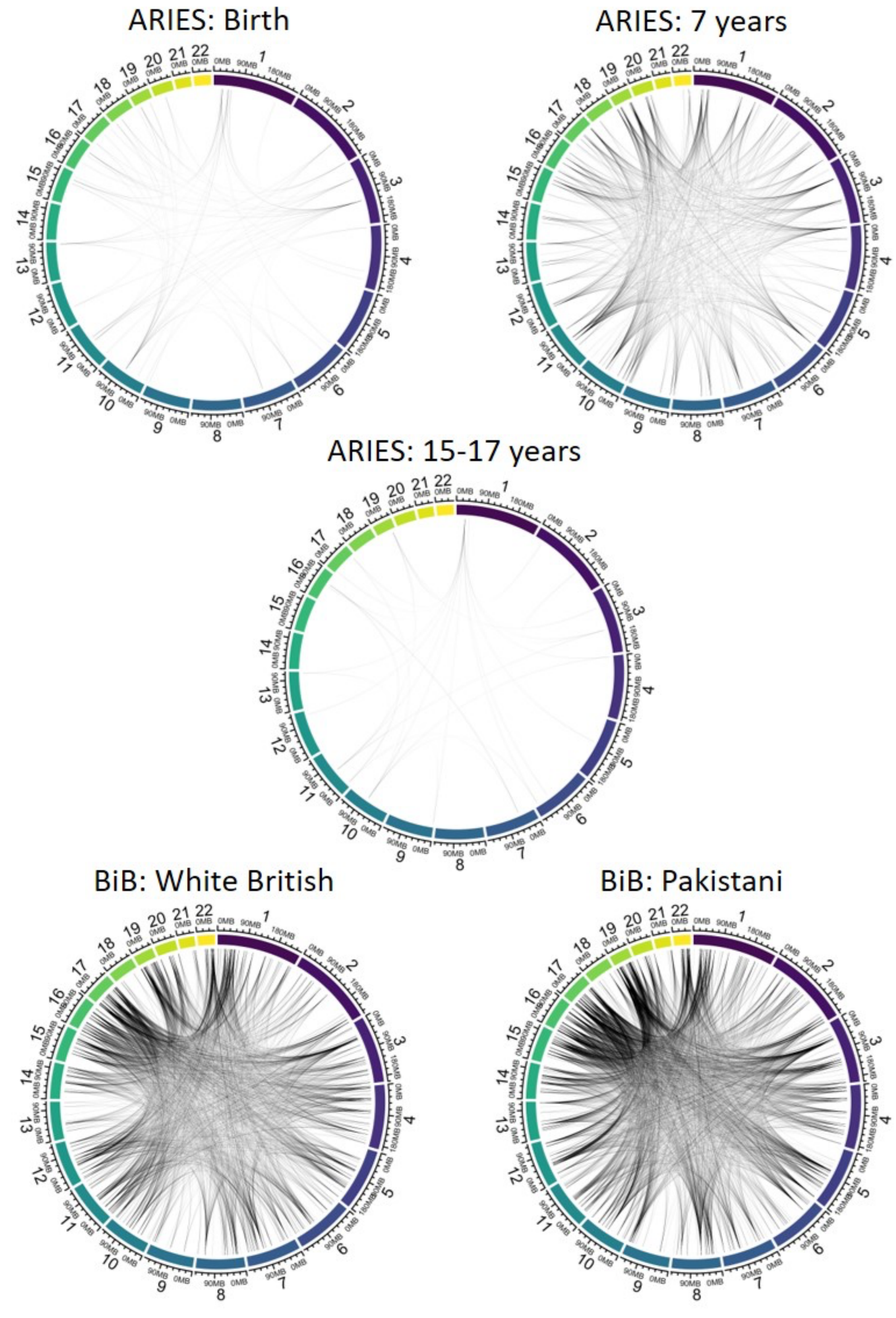

*Figure 8:* ***Circos plots*** *visualising trans correlations r>0.9 in ARIES at birth (top left), 7 years (top right) and 15-17 years (middle), and in BiB in the white British group (bottom left) and the Pakistani group (bottom right).*

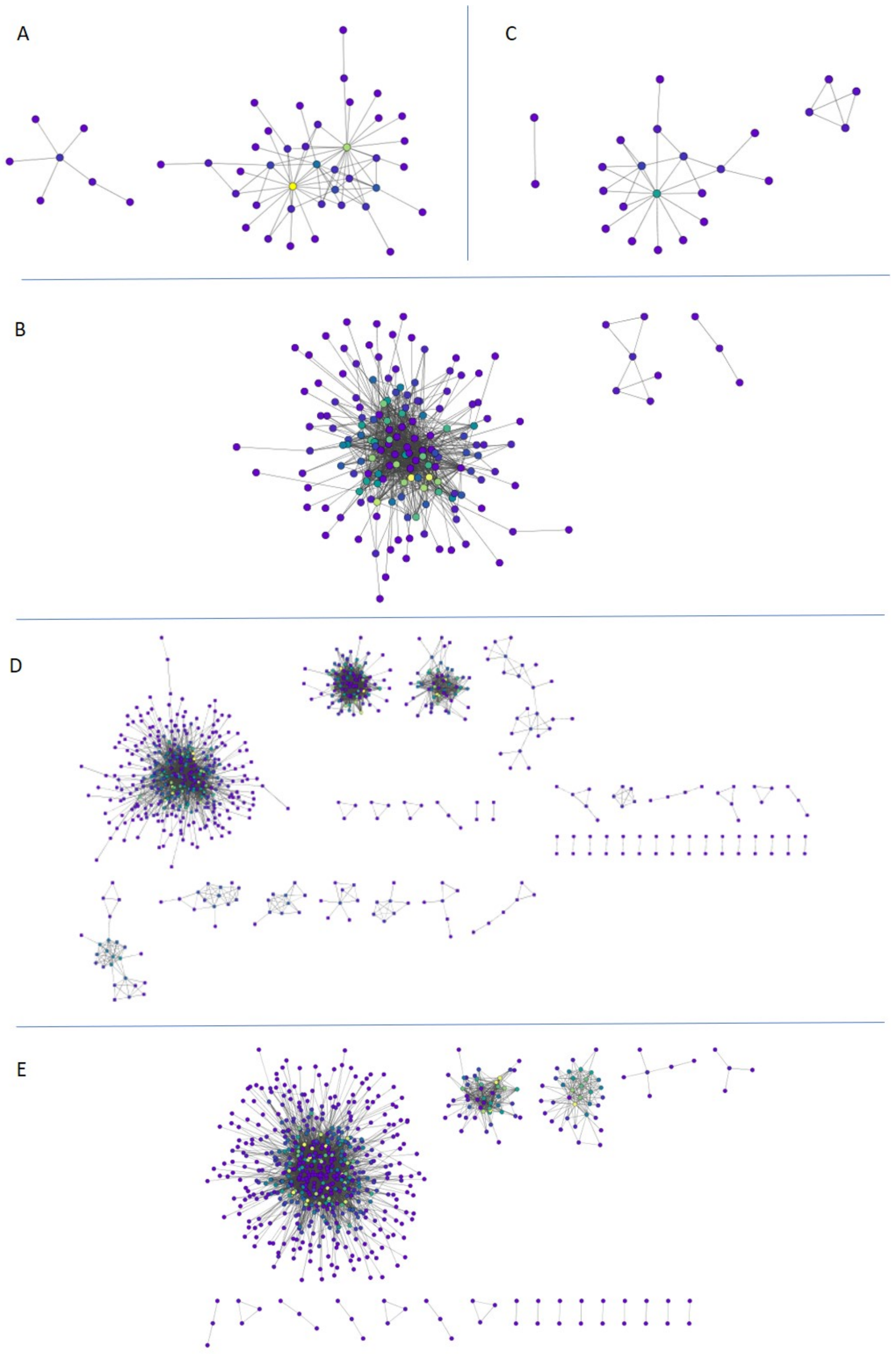

*Figure 9: **Cytoscape network plots** illustrating trans correlations r>0.9 in ARIES **A** at birth **B** at 7 years **C** at 15-17 years, and in BiB **D** White British and **E** Pakistani groups*

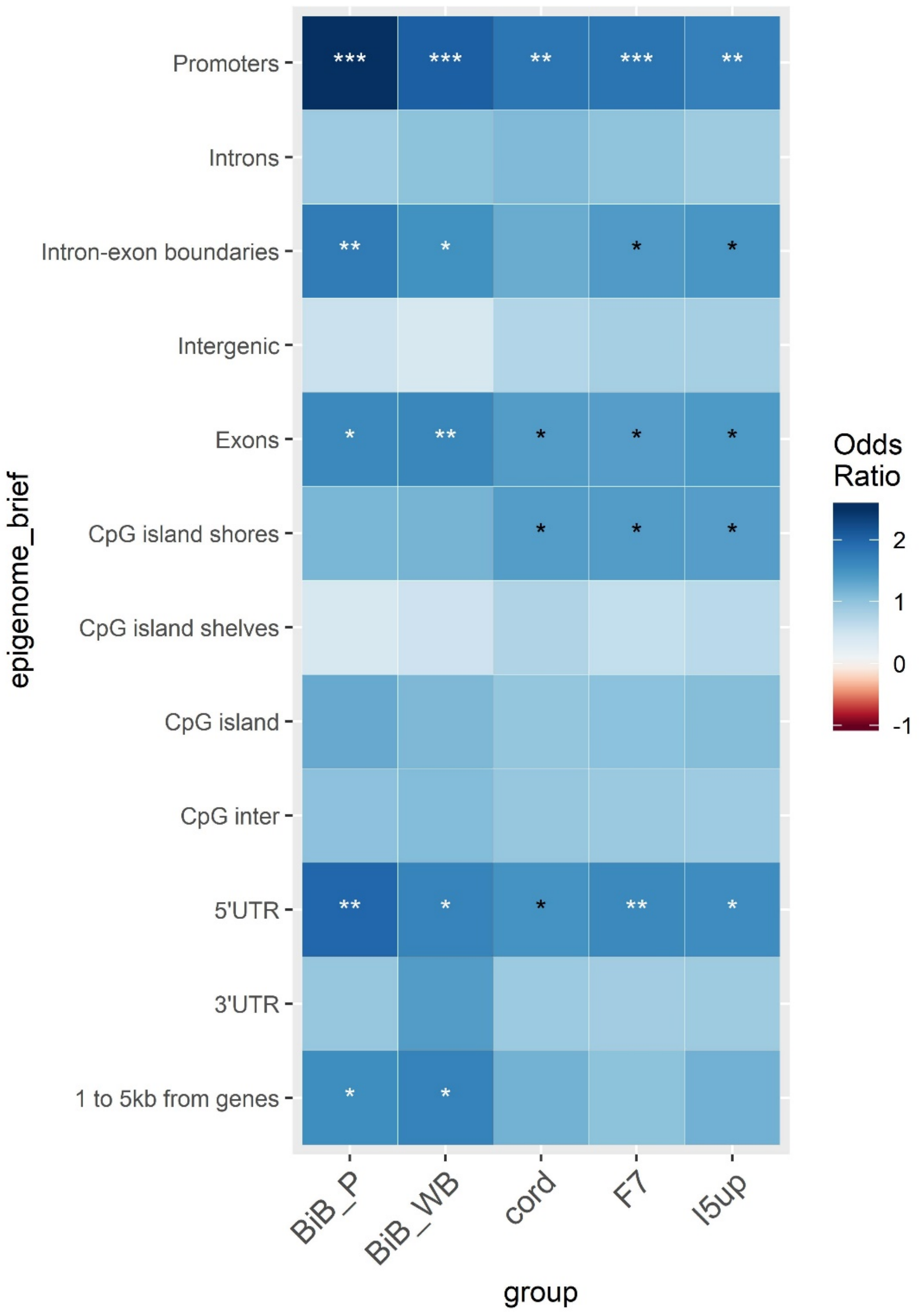

*Figure 10: Enrichment of genomic regions for cis correlating sites*

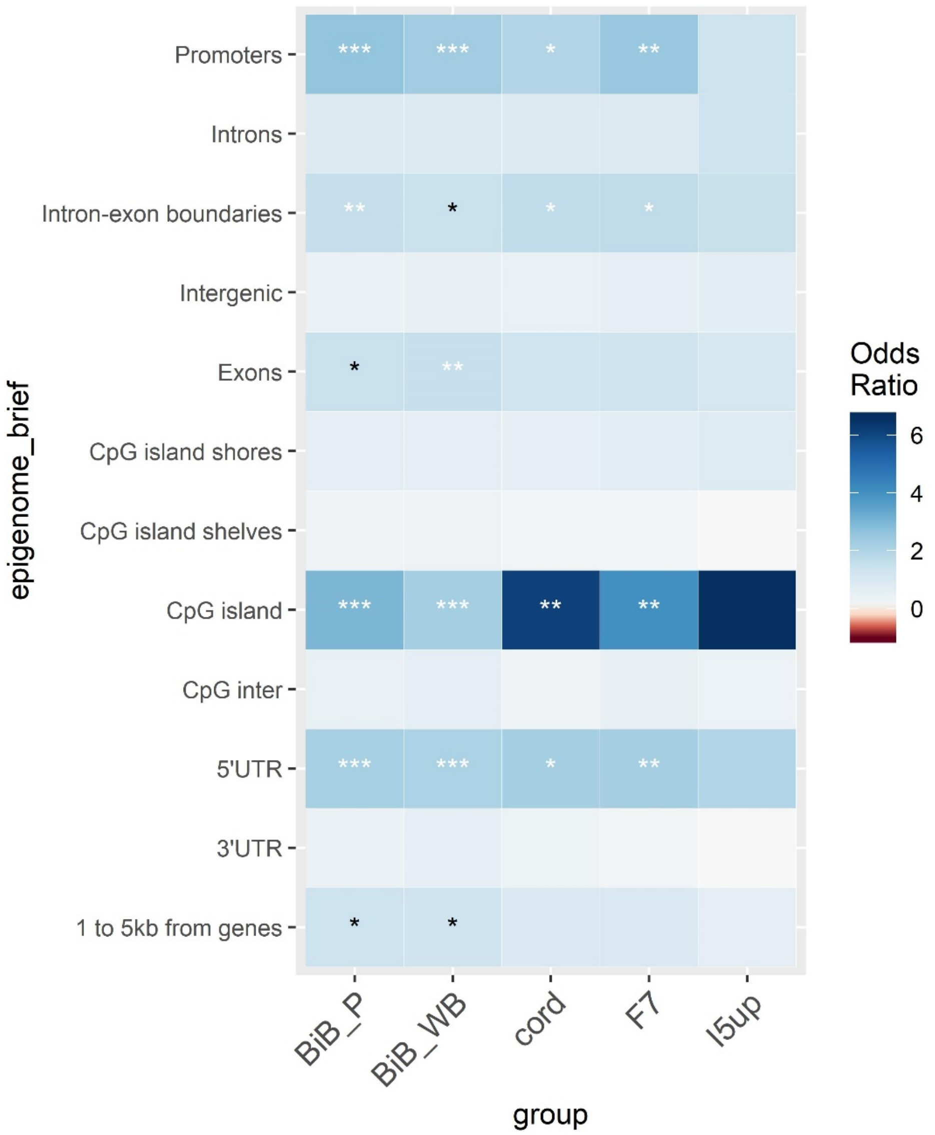

*Figure 11: Enrichment of genomic regions for trans correlating sites*
