## Supplementary methods for "DNA co-methylation has a stable structure and is related to specific aspects of genome regulation"

### Cohort descriptions

#### ARIES

The Avon Longitudinal Study of Parents and Children (ALSPAC) is a multi-generational cohort study based in Bristol. ALSPAC was set up with the aim to investigate factors that influence child health and development, and has collected data about a great number of exposures and outcomes related to this, including genetic and epigenetic data. All pregnant women in the former Avon region with a delivery date between April 1991 and December 1992 were eligible to take part in the study, and 14,541 pregnancies were initially recruited, with 14,062 live births (1, 2). Recruitment numbers were increased some years later, where individuals who were eligible at the start of the study but did not join were contacted (as long as they had not declined to take part in the study initially). 913 more children were recruited, giving a total of 15,454 pregnancies recruited, and 14,901 participants alive at one year old. Data has been collected frequently from ALSPAC participants, from pregnancy through to the present day, through questionnaires, clinics, biological samples and linkage data (1). The ALSPAC website contains a searchable data dictionary, containing all available data on the cohort, available at http://www.bristol.ac.uk/alspac/researchers/our-data/

Ethical approval for the study was obtained from the ALSPAC Ethics and Law Committee and the Local Research Ethics Committees. http://www.bristol.ac.uk/alspac/researchers/research-ethics/ proposal number B2808.

ARIES (Accessible Resource for Integrated Epigenomic Studies) is a sub-sample of the ALSPAC study, described in detail by (3). ARIES consists of 1022 mother-child pairs from ALSPAC, who were selected because they had appropriate DNA samples for DNAm profiling from specified timepoints. Participants included in ARIES are reasonably representative of those in ALSPAC as a whole; however ARIES mothers were slightly older, less likely to smoke during pregnancy, and more likely to have a non-manual occupation (3). As shown in Table 5 in the main text, ARIES participants were most likely to be in the high household class group. DNA methylation was profiled at two timepoints for the mothers: during pregnancy, and at follow-up approximately 15-17 years later. In the children, it was profiled at birth (using a sample of umbilical cord blood), during childhood (mean age 7.5) and during adolescence (mean age 17.1 years) (3). This resulted in 5469 DNAm profiles.

#### BiB

Born In Bradford (BiB) is a longitudinal, multi-ethnic cohort study based in Bradford, UK. Like ALSPAC it was set up to investigate factors which influence child health and development, but with a particular focus on child morbidity and mortality, as rates of these have been higher in Bradford than the rest of the UK (4). Bradford has a high rate of economic deprivation – one third of the neighbourhoods in Bradford are in the most deprived 10% of neighbourhoods in England (5, 6). Around 20% of the population are of South Asian descent (4), so this cohort also provides the opportunity to study correlation between DNAm sites in a non-European ethnic group.

All women booking an oral glucose tolerance test (OGTT) at the Bradford Royal Infirmary at 26-28 weeks' gestation (around 80% of expectant mothers in Bradford) were invited to take part in the study. At the OGTT appointment, consent was obtained from those who were willing to take part (about 80% of those attending), and those who did not attend the OGTT were contacted at other hospital appointments. In total 12,453 women with 13,776 pregnancies were recruited, leading to 13,740 live births. Ethical approval for this study was granted by the Bradford Research Ethics

Committee (Ref 07/H1302/112). Written informed consent was obtained from the mothers (for themselves and their children) when they registered for the study.

A subsample of 1000 mothers and their children in BiB had DNAm data generated from blood samples taken during pregnancy in the mothers, and from cord blood in the children. Eligibility for this subsample was defined as mothers who had both completed the OGTT, had genetic data available (which comprised around 65% of the individuals with the completed OGTT) and who had complete self-report race/ethnicity data provided as part of the questionnaire completed at recruitment. The questionnaire offered nine racial/ethnic groups for selection, of which all participants with DNAm data were either white British or Pakistani origin. This subsample was specifically designed to be multi-ethnic, and so of those eligible, 500 White British and 500 Pakistani mothers were selected to have DNAm generated for themselves and their children.

### DNA methylation data generation

#### ARIES

Consent for biological samples was collected in accordance with the Human Tissue Act (2004). Standard procedures were used to collect blood samples. At birth, all samples were cord blood. All other timepoints were whole blood or buffy coat. DNA was extracted from samples and bisulfite converted with the Zymo EZ DNA MethylationTM kit (Zymo, Irvine, CA). DNA methylation was then profiled using the Illumina Infinium HumanMethylation450K BeadChip, using the standard protocol. An Illumina iScan was used to scan the arrays, and GenomeStudio (version 2011.1) was used for initial data quality review (3). The Beta-value statistic was used to represent methylation levels. Low quality profiles were removed, leaving 4593 for further analysis. All ARIES samples were run at the same time, so samples from all timepoints were semi-randomised across Beadarrays (referred to from here as slide), to minimise the likelihood of batch effects inducing differences between timepoints. A semi-randomisation procedure was used rather than a fully random one, to ensure samples from different timepoints were distributed across the arrays. Before normalisation, 615 outlying samples were removed from the dataset. The details of outlier removal can be found in (7). 21 further individuals were removed from the full ARIES dataset as they were the only sample on a slide to avoid errors running regression in R.

#### BiB

To generate DNAm data, the EZ-96 DNA methylation kit (Zymo Research, Orange, CA, USA) was used to bisulfite-convert 500ng of high molecular weight DNA. DNAm was assessed using the Illumina Infinium MethylationEPIC beadchip arrays (Illumina, San Diego, CA, USA). Batch variables were recorded using a laboratory information management system (LIMS). The Beta-value statistic was used to represent methylation levels. 2010 samples (mothers, children and control samples) were generated together. 98 failed genotype concordance checks, and 13 failed other QC checks, and as some of these issues overlapped in samples from the same individuals, a total of 100 individuals were removed during QC. This left 1910 samples for normalisation, comprising 25 controls, 934 mother and 951 child participants. For this study we used DNAm data only for the children; 88 further participants were removed as they were related >12.5%.

### DNA methylation data normalisation

All ARIES timepoints were normalised together, as they were all run at the same time. Both BiB racial/ethnic groups were normalised together. Methylation data for both cohorts were normalised using the Functional Normalization algorithm (8) implemented in meffil, using the top 10 principal components from the control probes; full details of this have been published in the meffil paper (7). Slide effects were notable even after normalisation, so slide was regressed out from the raw methylation betas before normalisation. This was to remove some of the slide effects so that the technical artefacts could be better captured by the control probes. Slide row has also been shown to affect DNAm data; however this is picked up by the staining control probes, and was regressed out during the normalisation.

### Removing outlying methylation values

For each DNAm site observations that were more than 10 standard deviations from the mean were removed from the data, repeating this process three times to remove sufficient outliers. Where outliers were removed, they were replaced by the mean for that probe. There is no clear consensus about whether outliers should be removed from DNAm data although recent work suggests that outlying DNAm values represent rare genetic variants (9). However the aim of the present analysis is to aggregate DNAm measurements across nearly 1000 samples, and so outliers caused by rare genetic variants will simply serve to skew the estimates without contributing information useful to the present analysis. The outlying values were replaced with the mean for that probe for practical reasons, as missing values would cause problems in computing residuals and correlations. There were a small number of outliers in comparison to the size of the methylation matrix (around 100,000 per dataset, which is around 900 x 394,842); equivalent to approximately one quarter of the DNAm sites having a single outlying value.

### Filtering DNAm data

In ARIES the Zhou et al (10) list of probes was used to exclude sub-optimal DNAm sites from the analysis. Also removed were DNAm sites which were identified as multi-mapping probes (bisulfite converted sequences allowing two mismatches at any position mapped to the hg19 primary assembly, i.e. no haplotypes included as in (11), probes with variants (MAF >5%, UK10K) at the CpG dinucleotide or the extension base (for type I probes), and any probes targeting non CpG sites that failed liftover to hg19 (12). As BiB uses the EPIC array to measure DNAm, we reduced the BiB DNAm data to sites which were present in the ARIES dataset after filtering out sub-optimal sites.

### Genotype data generation

#### ARIES

ARIES participants were genotyped as part of the main ALSPAC study. All ALSPAC child participants were genotyped with the Illumina HumanHap550 quad genome-wide SNP array (Illumina Inc., San Diego, CA) by the Laboratory Corporation of America (LCA, Burlington, NC, USA) and the Wellcome Trust Sanger Institute (WTSI, Cambridge, UK), supported by 23andMe (3). Participants were excluded if they had the incorrect gender assigned, if there was abnormal heterozygosity (defined as <0.310 or >0.330 for the LCA data, and <0.320 or >0.345 for the WTSI data), high missingness (>3 %), if there was cryptic relatedness (>10 % identity by descent), and if the individual was of non-European ancestry (which was detected by multidimensional scaling analysis). After QC, the dataset consisted of 500,527 directly genotyped SNP loci. SNP data were then imputed to increase SNP density. They were imputed using the 1000 Genomes reference panel (phase 1, version 3, phased using SHAPEIT (version 2, December 2013) (13), using all populations (14)), using IMPUTE (v2.2.2) (15, 16). Genotypes were retained if they had Hardy Weinberg equilibrium $p > 5e{-7}$, a minor allele frequency of more than 1%, and an imputation info score over 0.8.

#### BiB

Samples in BiB were genotyped using either the Illumina HumanCoreExome Exome-24 v1.1 microarray, or the Infinium global screen-24+v1.0 array. GenomeStudio 2011.1 was used to pre-process samples. If samples had a call rate of <0.95, they were excluded. Poorly performing SNPs were removed. Most multi-allelic SNPs were discarded. 459,340 SNPs remained, and these were imputed by the Sanger Impute Service using the 1000genomes and UK10K reference panels (as 1000 genomes contains a number of different racial/ethnic groups).

### References


1. Boyd A, Golding J, Macleod J, Lawlor DA, Fraser A, Henderson J, et al. Cohort Profile: the 'children of the 90s'--the index offspring of the Avon Longitudinal Study of Parents and Children. Int J Epidemiol. 2013;42(1):111-27.
2. Fraser A, Macdonald-Wallis C, Tilling K, Boyd A, Golding J, Davey Smith G, et al. Cohort Profile: the Avon Longitudinal Study of Parents and Children: ALSPAC mothers cohort. Int J Epidemiol. 2013;42(1):97-110.
3. Relton CL, Gaunt T, McArdle W, Ho K, Duggirala A, Shihab H, et al. Data Resource Profile: Accessible Resource for Integrated Epigenomic Studies (ARIES). Int J Epidemiol. 2015;44(4):1181-90.
4. Wright J, Small N, Raynor P, Tuffnell D, Bhopal R, Cameron N, et al. Cohort Profile: the Born in Bradford multi-ethnic family cohort study. Int J Epidemiol. 2013;42(4):978-91.
5. Department for Communities and Local Government. The English Indices of Deprivation 2015: Statistical Release 2015 [Available from: https://assets.publishing.service.gov.uk/government/uploads/system/uploads/attachment_data/file/465791/English_Indices_of_Deprivation_2015_-_Statistical_Release.pdf.
6. Smith T, Noble M, Noble S, Wright G, McLennan D, Plunkett E. The English indices of deprivation 2015. London: Department for Communities and Local Government. 2015.
7. Min JL, Hemani G, Davey Smith G, Relton C, Suderman M. Meffil: efficient normalization and analysis of very large DNA methylation datasets. Bioinformatics. 2018;34(23):3983-9.
8. Fortin JP, Labbe A, Lemire M, Zanke BW, Hudson TJ, Fertig EJ, et al. Functional normalization of 450k methylation array data improves replication in large cancer studies. Genome Biol. 2014;15(12):503.
9. Chundru VK, Marioni RE, Pendergast JGD, Lin T, Beveridge AJ, Martin NG, et al. Rare Genetic Variants Underlie Outlying levels of DNA Methylation and Gene-Expression. bioRxiv. 2020.
10. Zhou W, Laird PW, Shen H. Comprehensive characterization, annotation and innovative use of Infinium DNA methylation BeadChip probes. Nucleic Acids Res. 2017;45(4):e22.
11. Naeem H, Wong NC, Chatterton Z, Hong MK, Pedersen JS, Corcoran NM, et al. Reducing the risk of false discovery enabling identification of biologically significant genome-wide methylation status using the HumanMethylation450 array. BMC Genomics. 2014;15:51.
12. Min JL, Hemani G, Hannon E, Dekkers KF, Castillo-Fernandez J, Luijk R, et al. Genomic and phenomic insights from an atlas of genetic effects on DNA methylation. medRxiv. 2020:2020.09.01.20180406.
13. Delaneau O, Marchini J, Genomes Project C, Genomes Project C. Integrating sequence and array data to create an improved 1000 Genomes Project haplotype reference panel. Nat Commun. 2014;5:3934.
14. Genomes Project C, Auton A, Brooks LD, Durbin RM, Garrison EP, Kang HM, et al. A global reference for human genetic variation. Nature. 2015;526(7571):68-74.
15. Howie B, Marchini J, Stephens M. Genotype imputation with thousands of genomes. G3 (Bethesda). 2011;1(6):457-70.
16. Howie BN, Donnelly P, Marchini J. A flexible and accurate genotype imputation method for the next generation of genome-wide association studies. PLoS Genet. 2009;5(6):e1000529.
